# The SARS-CoV-2 Spike Variant D614G Favors an Open Conformational State

**DOI:** 10.1101/2020.07.26.219741

**Authors:** Rachael A. Mansbach, Srirupa Chakraborty, Kien Nguyen, David C. Montefiori, Bette Korber, S. Gnanakaran

**Affiliations:** Theoretical Biology and Biophysics, Los Alamos National Laboratory, Los Alamos, NM 87545; Center for Nonlinear Studies, Los Alamos National Laboratory, Los Alamos, NM 87545; Duke Human Vaccine Institute & Department of Surgery, Durham, NC 27710

## Abstract

The COVID-19 pandemic underwent a rapid transition with the emergence of a SARS-CoV-2 variant that carried the amino acid substitution D614G in the Spike protein that became globally prevalent. The G-form is both more infectious *in vitro* and associated with increased viral loads in infected people. To gain insight into the mechanism underlying these distinctive characteristics, we employed multiple replicas of microsecond all-atom simulations to probe the molecular-level impact of this substitution on Spike’s closed and open states. The open state enables Spike interactions with its human cellular receptor, ACE2. Here we show that changes in the inter-protomer energetics due to the D614G substitution favor a higher population of infection-capable (open) states. The inter-protomer interactions between S1 and S2 subunits in the open state of the D-form are asymmetric. This asymmetry is resolved in the G-form due to the release of tensile hydrogen bonds resulting in an increased population of open conformations. Thus, the increased infectivity of the G-form is likely due to a higher rate of profitable binding encounters with the host receptor. It is also predicted to be more neutralization sensitive due to enhanced exposure of the receptor binding domain, a key target region for neutralizing antibodies.

## Introduction

COVID-19 is caused by the coronavirus SARS-CoV-2, and the pandemic is a global emergency. The virus infects human cells through the binding of the receptor binding domain (RBD) of its trimeric Spike glycoprotein (**Fig. 1**) to the human angiotensin-converting enzyme 2 (ACE2) receptor. Once bound, the S1 region (**Fig. 1A**, red) of the protein detaches (“sheds”), and the S2 region (**Fig 1A**, blue) triggers membrane fusion and mediates viral entry. In the “all-down” Spike conformation, all three RBDs of the protomers are in the closed orientation (**Fig.1A**). Binding of the RBD to ACE2 is thought to require the transition of one of the protomers from the closed to an open conformation (1), (2) (**Fig. 1B**). This infection-capable configuration is referred to as “one-up” Spike conformation. The “two up” and “three up” states are less stable since they are captured only by stabilizing mutations in SARS-CoV1 (3) or by introducing disulfide bonds in SARS-CoV2 (4).

**Fig. 1.**
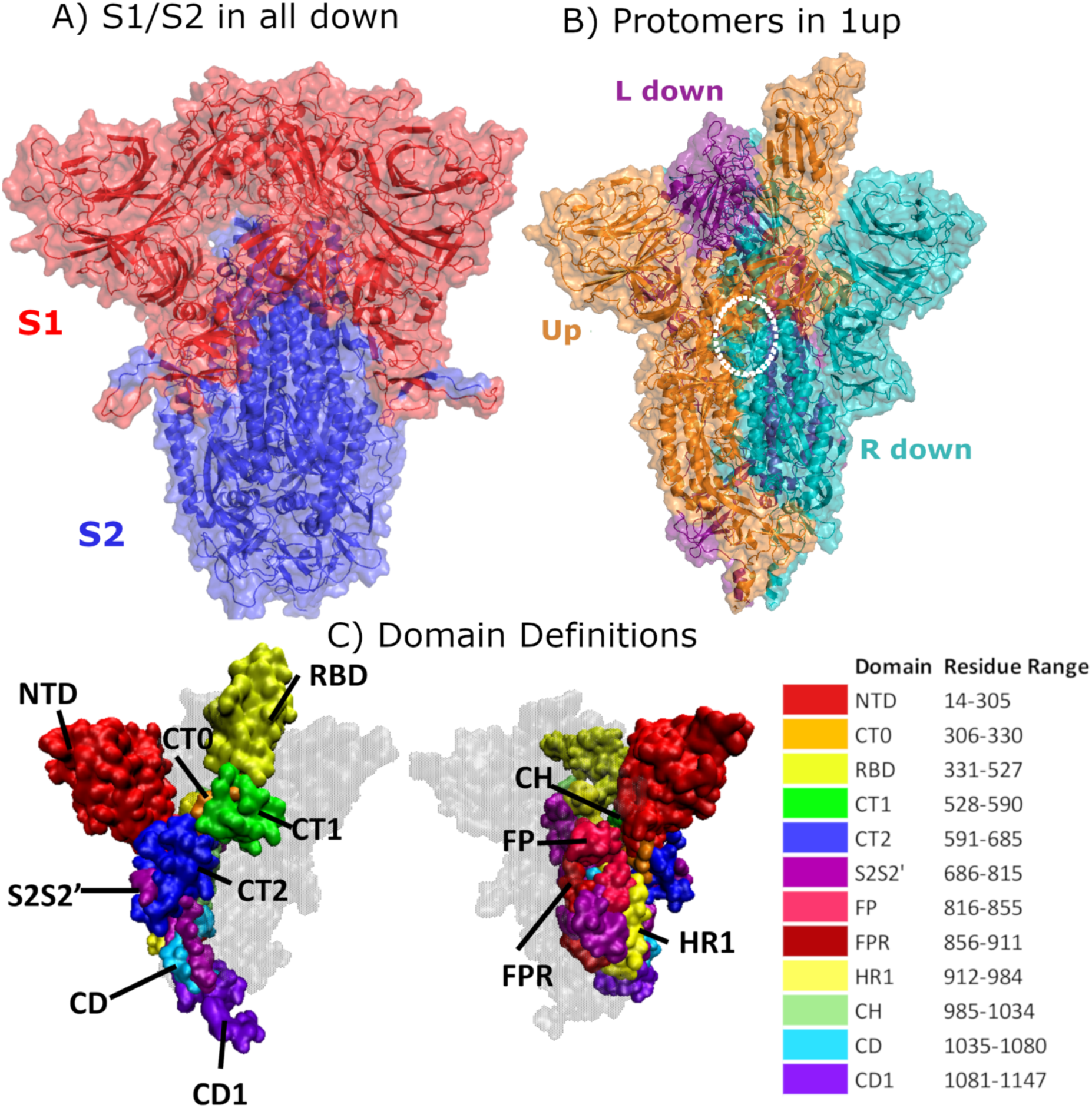
Structural Representation of the Spike protein. **(A)** The Spike complex is shown in the “all-down” conformation. Its S1 and S2 subunits are depicted in red and blue. **(B)** The Spike complex is shown in the “one-up” conformation. It is composed of three protomers, with the L-down protomer depicted in purple, the Up protomer in orange, and the R-down protomer in cyan. The dashed white circle indicates a D614G site. Here, only one of the three D614G sites are shown. **(C)** Definition of twelve domains and their residues used for analysis of simulations. It was necessary to define domains for every region for analysis, so some regions may have arbitrarily assigned names or not follow canonical sequence ranges. We display the domains highlighted in the Spike structure, shown from two different perspectives. Definitions of domain abbreviations: NTD, N-Terminal Domain; CT0, C-Terminal domain 0; RBD, Receptor Binding Domain; CT1, C-Terminal domain 1; CT2, C-Terminal domain 2; S2S2’, S1/S2 Cleavage to S2’; FP, Fusion Peptide; FPR, Fusion Peptide Region; HR1, Heptad Repeat 1; CH, Center Helix; CD, Connector Domain; and CD1, Connector Domain 1. Images in **(A)-(B)** were created with PyMol (15); images in **(C)** were prepared using VMD (16).

A viral variant has recently emerged, carrying a single amino acid substitution in Spike at residue 614 from an aspartic acid (D) to a glycine (G) (D614G). This G-form is now the globally prevalent form and is potentially more transmissible (5). It is associated with increased viral nucleic acid in the upper respiratory tract (5) and has higher infectivity in pseudotype virus assays in multiple cell types (5), (6). To identify the molecular factors driving the experimentally-observed differences, we performed cumulative 20 microsecond-length all-atom molecular dynamics (MD) simulations of the full Spike trimer in explicit solvent. We generated four separate sets of simulations, each with five replicas, to study both the D614 form and G614 forms, in either the all-down or one-up states. Using these simulations, we compared interactions formed between twelve regions of the Spike protein (**Fig. 1C)**. To identify the factors that lead to distinct dynamics for each of the four models, we compared residue-residue interactions and correlations, hydrogen bonding, and exposure of ACE-2 binding and RBD epitope sites.

We found that the D614G substitution was directly associated with far-reaching alterations in inter-protomer interactions and indirectly associated with changes in binding site exposure. Specifically, G614 form leads to a rearrangement in residue-residue contacts and hydrogen bonding that would lead to increased population of the one-up Spike ensemble. A careful analysis of RBD exposure, including the role of the highly-flexible glycans, showed that both the ACE2 binding site and critical neutralizing antibody epitopes in the RBD are more accessible in the one-up state (1), (7), (8).

## Results

Individual protomers of the Spike undergo distinct fluctuations (**Fig. S1**). To distinguish between protomers when discussing the results, we refer to the one that transitions between up and down orientations state as the “Up” protomer. The other two protomers that are in the down configurations are referred to as, “L-down” and “R-down”, depending whether the protomer is on the left or right of the Up protomer, respectively.

### Protein contacts are more symmetric between protomers in the G614 one-up state

To elucidate how specific interactions are altered by the D614G substitution, we calculated residue-residue contacts (9) that are formed between subunits of all three protomers. Specifically, these contacts include (a) S1-S2 interactions within a protomer (i.e. intra-protomer contacts), and (b) S1-S1, S2-S2, and S1-S2 interactions where the subunits are from different protomers (i.e. inter-protomer contacts). Two residues were considered to be in contact if their minimum distance was within a cutoff of 6Å. Contacts that occurred with probabilities of at least 50% were defined as “persistent” contacts.

The average number of persistent intra-protomer contacts between S1 and S2 was not affected by the D614G substitution in either the open or closed Spike conformation (**Fig. 2A-B**, first three data points). Similarly, regardless of the conformational state, the inter-protomer interactions between S1-S1 and between S2-S2 subunits were largely unaffected by the substitution (**Fig. S2A-B**). In contrast, the inter-protomer S1-S2 interactions are significantly impacted by the D614G substitution in the open conformation (**Fig. 2A-B**, last three datapoints). Specifically, in the all-down case, both the G- and D-forms lead to persistent numbers of contacts that are similar (within error bars) across all three protomer interfaces. This inter-protomer contact signature of the all-down Spike protein can be described as “symmetric.” In contrast, the D614 form loses the symmetrical contact signature when the Spike protein adopts the one-up conformation (**Fig. 2B, Table S1**), while the one-up G-form complex maintains the symmetry in the number of persistent contacts between the three protomers.

**Fig. 2.**
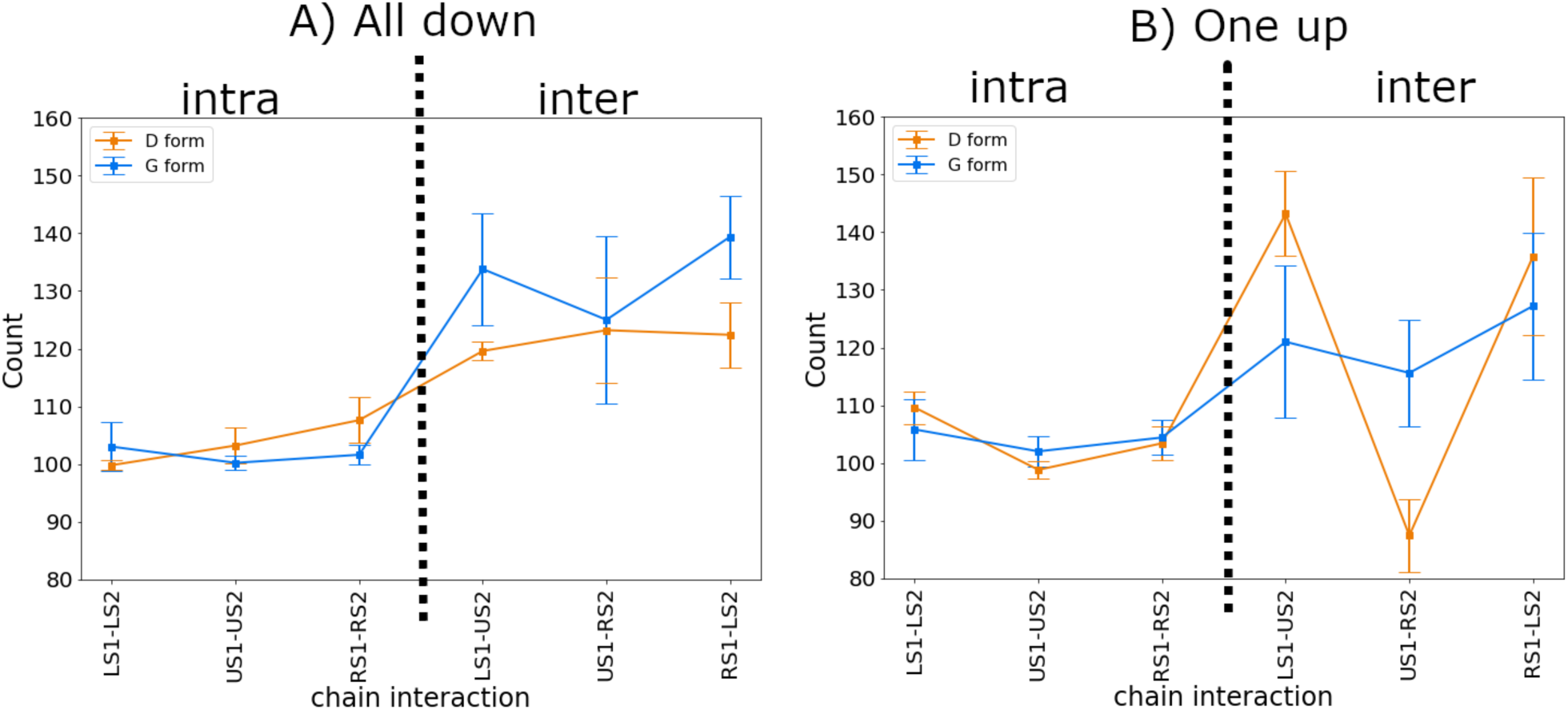
D614G substitution alters inter-protomer interactions between S1 and S2 subunits. **(A-B)**. Total number of contacts formed at S1-S2 interfaces **(A)** in the all-down system and **(B)** in the one-up system. Appropriate labelling and dashed lines indicate intra-protomer and inter-protomer S1-S2 interactions. For each set of simulations, error bars were calculated as standard error across five replicas. In the x-axis of all panels, ‘L’ denotes the L-down protomer, ‘U’ the Up protomer, and ‘R’ the R down protomer. For instance, LS1-U2 represents the interactions between the S1 region of the L-down protomer and the S2 region of the Up protomer.

### G-form RBD dynamics influenced by symmetric inter-protomer S1-S2 interactions

To further explore the factors that lead to the observed contact asymmetry, we calculated the cross-correlation matrix ***C*** of the Cα atoms for the entire complex (**Figs. 3, S3-S4)**. The greatest differences between the all-down state and the one-up state lie in the RBD-RBD and N terminal domain (NTD)-RBD correlations (**Fig. S3M-P**). Consistent with the contact analysis above, we observe greater symmetry in the G614 form one-up state, relative to the D614 form one-up state: the magnitude of the correlations between and within all three RBD domains are highly similar for the G614 form (**Fig. 3A**), but the intra-domain correlations and the U-RBD/R-RBD (anti)-correlations are stronger than the other inter-domain correlations in the D614 form (**Fig. 3B**, four squares in the lower right). There is also a slight asymmetry in the inter-domain scorrelations of the G-form all-down system (**Fig. S4A**, off-diagonal elements), compared to the D-form all-down system (**Fig. S4B**, off-diagonal elements). It is consistent with a subtle deviation from symmetry observed for the all-down G-form (**Fig. 2A, Table S1**). Overall, atomistic correlations indicate that in the one-up state the motions of the RBDs in the G614 form are more synchronized than the D614 form. Such significant positional correlations distal to the 614 substitution site are a signature of allostery (10), (11).

**Fig. 3.**
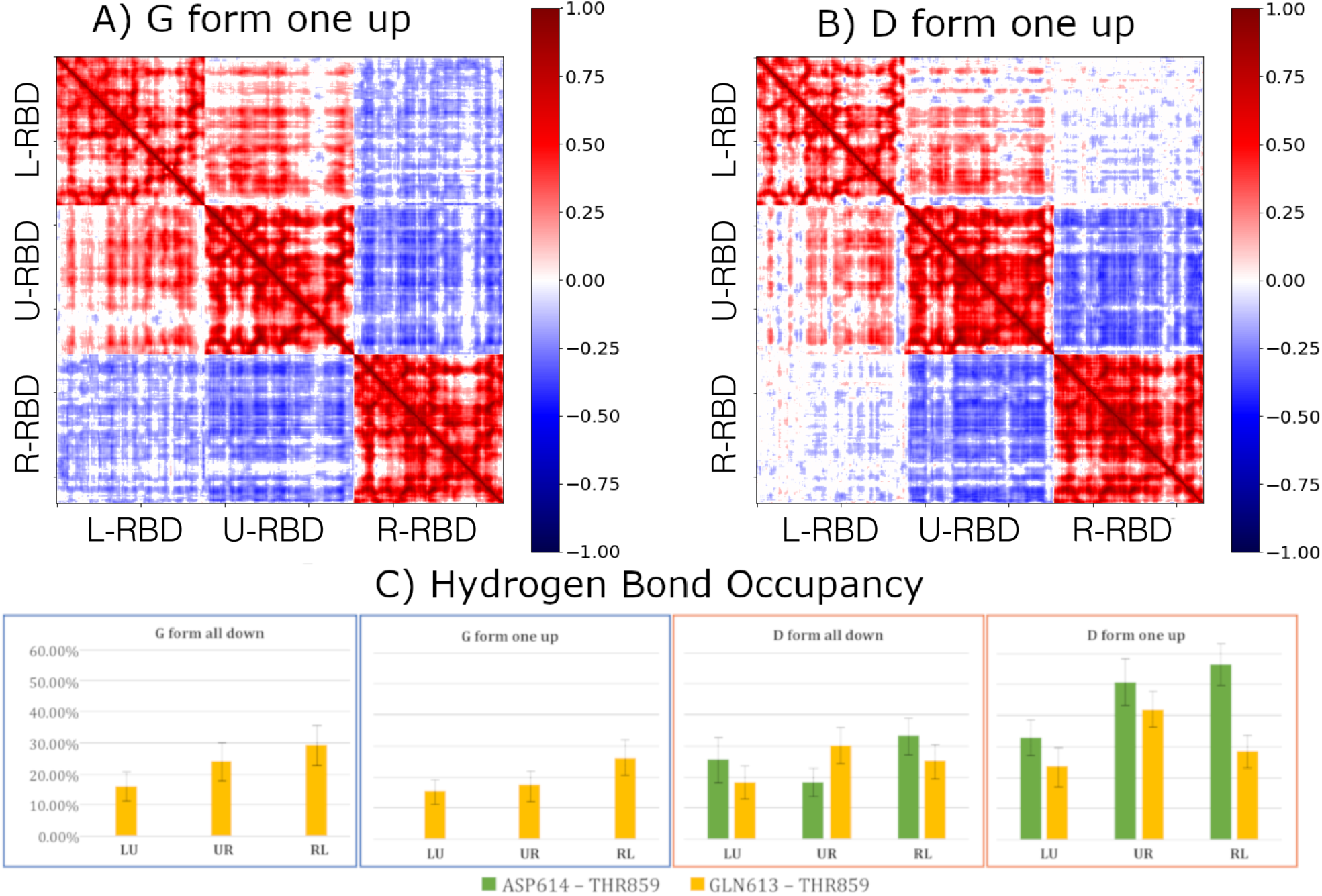
Residue-residue cross correlation matrices and hydrogen bonding. **(A)** G614 form one-up **(B)** D614 form one-up, In the x and y axes, L-RBD, U-RBD, and R-RBD denote RBD regions of the L-down, Up, and R-down protomers. **(C)** Hydrogen bond occupancy for critical residue pairs located between protomers. Occupancies were calculated for the D/G 614-THR59 pair and the GLN613-THR859 pair. For each system, error bars were calculated as standard error over five replicas. In all subpanels, ‘L’ represents the L-down protomer, ‘U’ the Up protomer, and ‘R’ the R-down protomer. For example, “LU” denotes the bonding between L and U protomers.

### Contacts in CT1-CT2 (528-685) and FP-FPR (816-911) regions are major contributors to the symmetrization in the G614 form

To capture the specific regions where the inter-protomer S1-S2 contacts are most affected by the D614G substitution, we carried out a global differential contact analysis, in which we identified persistent contacts that existed in one form but not the other in the all-down or one-up states. Although the overall number of contacts does not change for most of the interfaces, specific contacts do, with different contacts being gained at different interfaces (**Figs S5-S8**). Furthermore, the equalization of inter-protomer S1-S2 contacts in the one-up G614 form arises largely from the overall loss of contacts between the CT1-FP domains of L-down and Up and the gain of contacts between the CT1-FP and CT2-FP domains of Up and R-down (**Fig. S5**). Interestingly, the mutation of residues in the CT1, FP, and FPR domains—next to the FP domain—to more hydrophobic residues was experimentally linked to an increase of protomers adopting the up conformation. This provides experimental support for the importance of these regions in determining the conformational state of the Spike (4).

We next focused on studying the hydrogen bonding interactions of the speculated critical residues D/G 614 with T859 (5). The occupancy of the D614 – T859 hydrogen bond increases for the one-up versus all-down state in the D614 form (**Fig. 3C**, right two panels), but because the hydrogen bond no longer exists in the G614 form (**Fig. 3C**, left two panels), this bias likewise does not exist. This is another type of “symmetrization”. In addition, there is a concomitant increase in the flexibility of residues 610 to 650 in the G614 form (**Fig. S9G-J**). In general (**Figs S9-S10**), we do not observe compensatory hydrogen bond formation with THR 859. Thus, it is conceivable that this hydrogen bond reduction leads to a conformational relaxation.

### Symmetrization of inter-protomer interactions in G614 form can lead to higher population of infection-capable (one-up) conformation

The distinct contact signatures associated with the D614G substitution may be associated with alterations in the relative population of the all-down and one-up ensembles of the Spike protein (**Fig. 2**). For the G-form, the symmetry in the number of contacts among the inter-protomer interfaces suggests that the energetics between the protomers are similar. Since the G-form preserves these interactions in both the all-down and one-up conformations, the interaction energy between up- and down-protomers would be similar to the energetics between two down-protomers. Inspired by this rationale, we use an Ising model (12) to demonstrate how the D614G substitution can affect the relative stability of the open and closed Spike conformations. In this model, each protomer can adopt an “up” or “down” state, and each protomer interacts with its two neighbors with an energy that depends only on the states of both protomers. Defining the Spike as a periodic three-spin system, the Ising model suggests that symmetrization of the energetics of the G-form will lead to an ensemble ratio of 75:25 of the one-up state to the all-down state, as compared to the 50:50 population ratio for the original D-form (see SI for details). This demonstrates how a single residue substitution can facilitate the adoption of the open Spike configuration, thereby enhancing viral infection.

In summary, we find three contributions to symmetrizations associated with the D614G substitution: (1) symmetrization in the inter-protomer contacts, (2) symmetrization in the correlations between the RBDs, and (3) symmetrization of a specific inter-protomer hydrogen bond. We hypothesize that these three symmetrizations lead to a decrease in the energy differences between the all-down and one-up states, leading to an increase in infectivity through an increase in the one-up population of the G-form.

### One-up state enhances exposure of RBD to ACE2 binding and epitope sites

Since Spike is a glycoprotein, we also modeled the impact of highly-flexible glycan ensembles on RBD exposure and epitope shielding using a quantitative measure called the glycan encounter factor (GEF) (13). A low GEF value indicates a high exposure of the protein surface in a given region. We perform GEF analysis here to assess changes in exposure between all-down and one-up states and between the D- and G-forms (for details on local changes see **Supplementary Material** and **Fig. S11**). Independent of D- or G-forms, the ACE2 binding site and epitope regions are significantly more exposed in the one-up state than in the all-down state (**Fig. 4A-B)**. Specifically, the exposure increases by ∼40% at the Receptor Binding Motif (residues 438-506). This is accompanied by larger exposure of both the ACE2 binding site and RBD targeting antibody epitopes. For example, there is a ∼64% increase in exposure of the C105 epitope region (7) when the RBD adopts the open state, as opposed to the closed orientation where it is buried by surrounding protein and glycans (**Fig. 4C-E**). Thus, in the one-up state, which is favored by the G614 form, key epitopes of the RBD become more accessible compared to the D614 form.

**Fig. 4.**
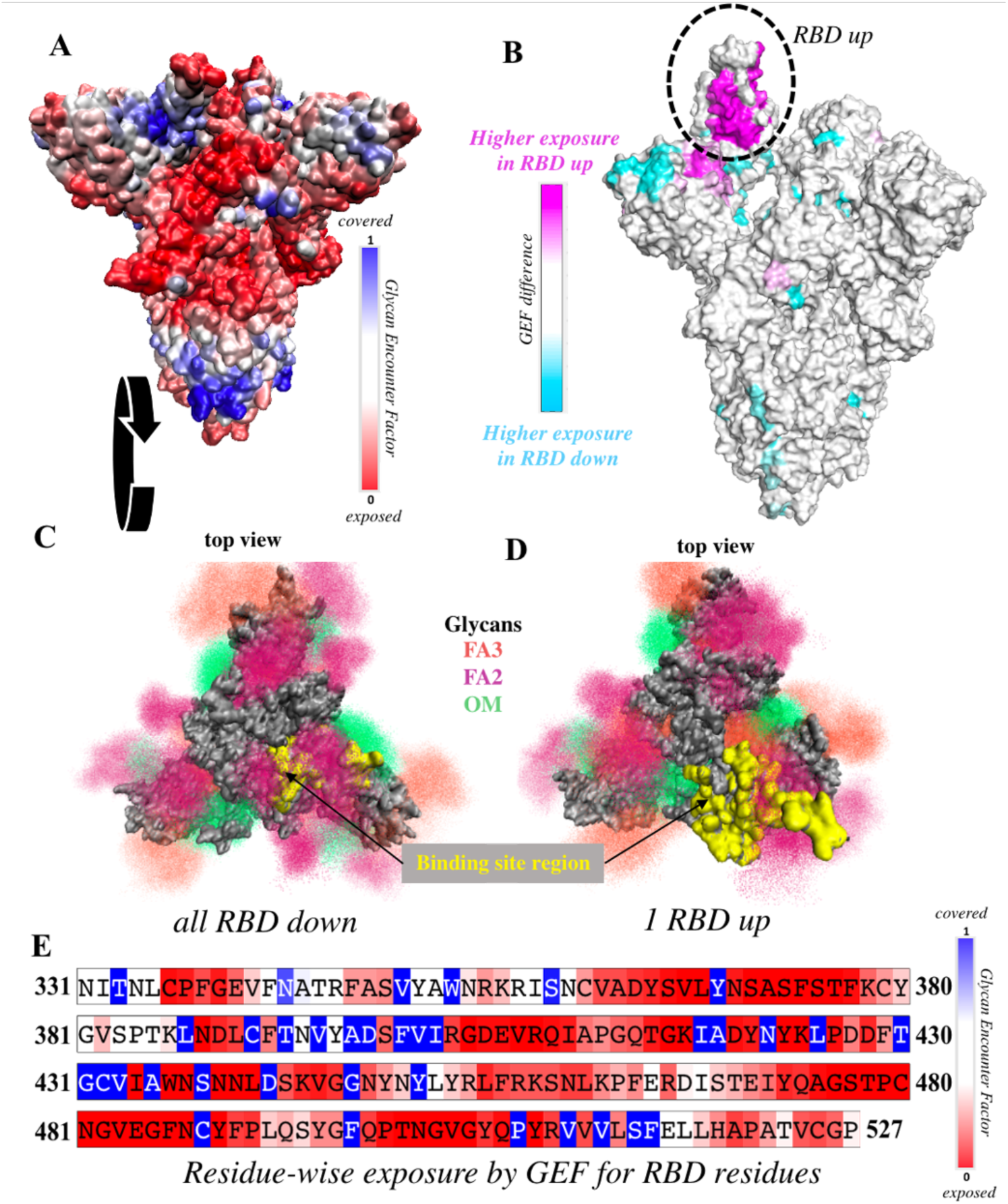
RBD is significantly more exposed in the one-up conformation. **(A)** Surface representation of the all-down Spike complex. Colors ranging between red, white and blue represent values for the glycan encounter factor (GEF). GEF describes relative exposure and coverage of protein surfaces. **(B**) Surface representation of the one-up Spike. Colors ranging between cyan, white, and magenta indicate GEF differences between the closed and open structures. The RBD in the up orientation is indicated by the dotted circle, where the originally buried (magenta) region around 458-478 is now exposed. **(C-D)** Top view of the Spike protein apex in **(C)** all-down and **(D**) one-up conformations. Protein surface is shown in grey. Glycan ensembles covering the protein surface are represented by point densities. Fucosylated 2 and 3 antennae complex (FA2 and FA3) glycans and oligomannose (OM) glycans are depicted in dark maroon, orange, and green. Binding site region, which includes receptor binding motif (residues 438-506) and C105 antibody binding residues (i.e. 403, 405, 406, 408, 409, 415-417, 420, 421, 449, 453, 455-460, 473-477, 486, 487, 489, 493-496, 498, 500-505), is colored yellow. These binding site residues are significantly more exposed in the one-up conformation, compared to the all-down case. **(E)** Residue-wise GEF of the RBD domain. Buried residues are colored blue. Exposure increases as the color changes toward white and red.

## Discussion

The D614G substitution occurs on the inter-protomer interface of the Spike protein mediating critical contacts. We have performed extensive all-atom MD simulations of the trimeric Spike protein and assessed the effects of this substitution on both all-down and one-up conformations. In parallel, Weissman and colleagues have obtained structural evidence using negative stain electron microscopy showing that the G614 form Spike populates the one-up state 82% of the time, while the D614 form adopts this open conformation with only 42% frequency. Further, they found that the G614 form is more sensitive to neutralization by sera raised from D614 form vaccinations in animal models (14). Our simulations provide mechanistic details that can explain the increased occupancy of the one-up state in the G614 form, which in turn accounts for both its enhanced infectivity (5), (6) and neutralization sensitivity (14).

We found that the different interactions between protomers in the D614 form are different in the one-up and all-down states. By contrast, in the G614 form there is a symmetrization in the number of inter-protomer contacts between S1 and S2 subunits, which is associated with allosteric fluctuations in RBD, distal to the 614 site. Thermodynamically, the effect of symmetrization can be captured using a periodic three-spin Ising model. Using the observed 50:50 ratio of all-down to one-up conformations in the D614 form (4), we show that symmetrization of the G-form energetics in comparison to the D-form would lead to a 75:25 ratio of one-up to all-down. It is noted that this ratio will become higher if there is a concomitant deviation from symmetry in the all-down state of the G614 form. The shift towards one-up Spike trimers would increase the likelihood of binding events of RBD and ACE2, thus explaining the experimentally observed increase in infectivity (5) and sensitivity to neutralization activity in vaccine sera (14).

Our dynamic model also indicates that there is a transition in the glycan shield when going from all-down to one-up; the glycan coverage disappears at the apex of the trimer when in the one-up conformation. Thus, regardless of the D- or G-form, the one-up conformation exposes the RBD for binding to the ACE2 receptor while simultaneously exposing more of the RBD protein surface for antibody binding to RBD epitopes. There are no significant global changes coming from the D614G substitution on the ACE2 binding site exposure in RBD once the Spike trimer is in the one-up conformation. Thus, given the natural preference for a more open Spike conformation in the G-form, it is possible that this form may have advantages as a vaccine antigen.

In summary, our findings suggest that the overall protein exposure remains globally similar between D- and G-forms, but predicts a dramatic increase in the one-up state with the G-form, meaning that both ACE2 binding and RBD-targeting antibody binding are likely to increase in the one-up state. Therefore, a change towards a higher one-up state population is likely the dominant effect of the D614G substitution. The mechanistic studies presented here, the structural data (14), the experimentally determined increase in infectivity (5), and the enhanced neutralizing antibody sensitivity (14) all come together in a consistent story.

## General

We thank Drew Weissman for sharing unpublished neutralization data and being willing to consider co-submission of our papers. We thank Priyamvada Acharya for sharing unpublished negative stain EM data on G614. We are grateful to Paul Weber for securing extensive computational resources that made this study possible.

## Author contributions

R.A.M., S.C., K.N., and S.G. designed the study. S.C. and K.N. prepared the structures. R.A.M. ran the simulations. S.C. calculated the GEF. S.C., K.N., R.A.M, B.K., D.C.M. and S.G. performed data analysis and interpretation. R.A.M and S.C. prepared the figures. R.A.M. wrote the initial manuscript draft. R.A.M., S.C., K.N., B.K., D.C. M., and S.G. rewrote and edited the manuscript. B.K. and S.G. secured the funding.

## Data and materials availability

Data and materials are available from the corresponding author upon reasonable request.

## Methods

### Structure Preparation

Atomic structures derived from cryo-EM (PDB IDs: 6VXX and 6VYB; Walls et al. (1)) were used to prepare the all-down and one-up configurations for simulation. As described in Ref (1), residues 986—987 were kept as Proline-Proline (PP), and Fusion Peptide residues 682—685 as SGAG instead of the viral form residues KV and RRAR, to maintain the stable soluble form of the spike extracellular domain protein. In the original 6VXX and 6VYB models, several flexible regions were unresolved, which vary in lengths from 2-27 residues. We used a data-driven structure-based modeling approach to build a complete model that accurately captures secondary structures of these missing regions, without introducing artificial ‘knotting’ of loops. For this, we employed homology modeling using numerous SARS-COV1 structures as templates (see **Supplementary Information**). Missing residues in the RBD were built using an ACE2-bound SARS-CoV2 structure (PDB: 6M0J) (2). Structure-based sequence alignment was performed using the 3D-Coffee program (3) and homology modeling was performed using the MODELLER 9.20 suite (4). We generated 10 models for each configuration and selected the top model based on DOPE and Procheck scores (5), (6). After testing that CryoEM fitting did not significantly improve the model employing Chimera 1.13.0 (7) rigid fitting, and the MDFF (8) flexible fitting, we arbitrarily chose as starting structures the all down model after 2 ns in vacuum with gscale = 0.3 and a further 2 ns in vacuum with gscale = 0.2 and the one up model after 2 ns in vacuum with gscale = 0.3. To avoid artificial charges at the protein ends, we introduced N-terminal acetylated and C-terminal N-methylamide capping groups. All histidines were modeled as the neutral tautomer where the epsilon nitrogen is protonated. 15 disulfide bonds found in 6VXX and 6VYB were modeled in the forcefield for simulation.

### Simulation Details

All-atom explicit-solvent simulations were performed with the Gromacs v5.1.2 and 2018.3 software packages (9), (10), using the CHARMM36m (11) forcefield and TIP3P water model (12). Each configuration was solvated and centered in a cubic box. The side length of the box was defined such that there is at least 15Å padding around the molecule. Each system was neutralized with an excess of 150mM KCL. Energy minimization was conducted using the steepest descent algorithm. Equilibration simulations were performed under constant number-volume-temperature (NVT, 2ns) and constant number-pressure-temperature (NPT, 10ns) ensembles. During the equilibration stages, harmonic position restraints were imposed on all heavy atoms of the molecule. Temperature coupling was achieved using Langevin dynamics at 310K with a relaxation time of 1ps (13). The Berendsen barostat with isotropic coupling was employed to maintain a constant pressure of 1 bar, with a relaxation time of 4ps and compressibility of 4.5×10E-5 / bar (14). Covalent bonds were constrained by implementing the LINCS algorithm (15). Van der Waals interactions were evaluated using a cutoff where forces smoothly switch to zero between 1.0 and 1.2nm. Coulomb interactions were calculated using the particle mesh Ewald (PME) method, with a cutoff of 1.2nm, a Fourier spacing of 0.12nm, an interpolation order of 4, and a tolerance of 1×10E-5 (16). Unrestrained production simulations were performed in the NPT ensemble, with an integration time step of 2fs. For each configuration, we simulated five replicas. Each replica was run for 1.1 µs. The first 100 ns of each run was considered as further equilibration and was not included for analysis. In total, we generated 20 1-µs production simulations.

### Contact analysis

We used the g_contacts plugin for Gromacs (17) to identify contact frequencies of all residue-residue contacts formed, both intra- and inter-protomer. A contact was defined as any heavy atom of one residue coming within 6Å of the heavy atom of another residue. For the contact analysis, we defined *persistent* contacts as any of the identified contacts that appeared with a frequency of greater than or equal to 0.5 – that is, that appeared in at least 50% of analyzed simulation frames. Average contacts with region S2 excluded the CD1 domain because of observations that it “frayed” in simulation and was extremely flexible, which we presume to be an artefact due to the lack of embedding of the Spike in the viral membrane.

### Correlation analysis

The cross-correlation matrix is defined as

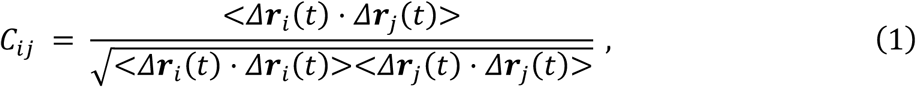

which is the normalized covariance where *Δ****r***_*i*_(*t*) is the fluctuations of atom *i* with respect to its average coordinates. We used the -cov, -norm, and -dot flags of the program carma to perform the analysis (18).

### Hydrogen bond analysis

We performed the hydrogen bond analysis with VMD 1.9.3 Hbonds plugin (19), where the definition of a hydrogen bond was the requirement of donor-acceptor atoms being polar, distance cutoff of 4Å and angle cutoff of 30^°^. This is a significantly more stringent criterion than that employed for the contact identification. A cutoff of 10% occupancy was used to identify extant hydrogen bonds.

### Glycan modeling and exposure calculation

There are 22 potential N-glycosylation sites (PNGS) in the Spike, each of which has a unique glycoform distribution. We chose glycan types with the highest occurrence at each site, as determined by mass spectrometry studies (20) for 19 of these glycans present within our model range (see **Fig. S11A** and **Table S3**). There are two putative O-glycan sites on the Spike; however, they have been reported to have less than 2% occupancy (21) and were therefore not included in our models.

We performed a Jarvis Patrick based cluster analysis where a structure is added to a cluster if it has at least 6 neighbors in common with another neighbor. Clustering was performed on all five replicas, for each of the four different systems, using the “gmx cluster” command with a cutoff of 0.2 nm to identify a subset of distinct conformations based on Cα RMSDs. We identified 63 clusters for D614 all-down, 78 clusters for D614 one-up, 54 clusters for D614G all-down, and 103 clusters for D614G one-up. We took the closest to the mean structure from each of the top twenty largest clusters of each system, and used these snapshots as the basis for building glycan models. Glycan ensemble modeling was performed using a previously established pipeline (22) with the ALLOSMOD (23) suite of MODELLER. An initial glycan conformation was added at each PNGS with randomized atomistic deviations from CHARMM36 ideal carbohydrate geometries, and random orientations. This was followed by simulated annealing relaxation of the entire glycoprotein. 50 glycosylated conformations were built on each of the 20 selected models, resulting in an ensemble of 1000 distinct glycoprotein conformations for each of the four systems. Glycoform for each PNGS was chosen as the one with the highest probability of occurrence from site-specific mass spectrometry results (20).

The Glycan Encounter Factor (GEF) was calculated for each residue exposed on the surface of the protein as established in our previous study (22). It is defined as the number of glycan heavy atoms encountered by an approaching probe of 6Å diameter mimicking a typical hairpin loop of antibodies interacting with epitopes. Geometric mean of this value measured in the three cardinal directions (perpendicular to the surface, and along the plane) was taken to cover different orientations.

### Limitations of the study

Although the results from these extensive MD simulations represent an extensive investment of computational resources, large-scale conformational shifts in the dynamics, such as the actual transition between the all-down and one-up states, are beyond the microsecond timescales considered here. Nonetheless, as long as they are applied on the appropriate timescale within the known limitations of the MD force field, the results of this article are of great significance in terms of explaining the molecular-level changes that occur as a result as a result of the D614G amino acid shift and its effects on the stability of different conformational states. Also, the current structure does not include the heptad repeat 2, trans-membrane or cytoplasmic regions that could differentially alter the presentation of Spike on the membrane in the G-form. We cannot comment directly on the S1/S2 cleavage site conformation or the effect of two prolines used for stabilization, as we used the sequence from Walls et. al (1) that was mutated to stabilize a soluble protein.

## Supplementary Materials

### Details of data-driven construction of initial structure via homology modeling

Soluble form of the SARS-CoV-2 Envelope spike protein was modeled based on available PDB structures. Structure 6VXX was used as the base template for all RBD down, closed model and 6VYB was used for the one-up model (1). These cryo-EM structures have several missing residues around the flexible loops. Ten such segments are more than 4 residues long, some are missing up to 29 residues. Analyzing the sequence with secondary-structure prediction-software JPRED-4 (2) and iTASSER (3), shows that many of them may have stabilized secondary structure features of helices and beta strands. Moreover, modeling of such long loops *ab initio* may lead to artifacts and ‘self-knotting’. Hence, we utilized available CoV-1 structure data as starting template to fill in these gaps whenever possible. Templates used for different loops are provided in **Table S2**.

Structure driven sequence alignment between CoV-1 and CoV-2 was used to determine template ranges, and is provided in **Fig. S12**. Following template selection, homology modeling was performed using a Variable Target Function Method of refinement, and 300 steps of template-restrained MD, with MODELLER 9.20 suite (4). 10 models were generated for each of the four systems. Model selection was based on DOPE score and PROCHECK stereochemistry scores (5). 15 disulfide bonds between cysteines pairs were identified and restrained with patches during modeling and appropriate force terms for these bonds were included for simulations. These are: 15-136, 131-166, 291-301, 336-361, 379-432, 391-525, 480-488, 538-590, 617-649, 662-671, 738-760, 743-749, 840-851.

### RMSD and RMSF for the four systems

Root mean square deviation (RMSD) at time *t* is calculated as,

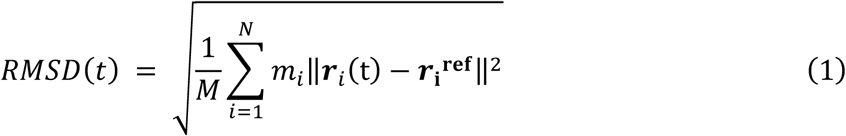

where *N* is the total number of atoms, 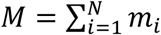is the total mass, 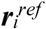 is the coordinates of the reference structure, and ***r*** *i* is the coordinates of the *i* th frame. All calculations were performed with the gmx rms tool from Gromacs 5.1.2 (6).

Root mean square fluctuation (RMSF) of atom *i* is calculated as,

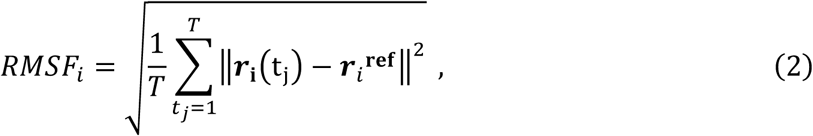

where *T* is the total time of the production simulation.

### Fluctuations of the one-up state

To describe RBD fluctuations relative to the rest of the protein, we computed the root mean square deviation (RMSD) of the RBD of each protomer from the initial equilibrated frame of each simulation after translational and rotational fit of the domains other than the RBD and the highly-flexible CD1. We also computed the root mean square fluctuations (RMSF) of each atom after translational and rotational fit of the Cα atoms of the whole protein. In **Fig. S1A**,**C**,**E**,**G**, we present the RMSD distributions of the RBDs. On the whole, the RMSDs of the three protomers are overlapping in the all-down configuration, whereas in the one-up conformation the L-down protomer shows smaller deviations. Specifically, in the all-down state, the overall means and standard deviations are *RMSD* _*L*_ = 0.7, σ_*RMSD, L*_ = 0.1, *RMSD* _*UP*_ = 1.0, σ*_RMSD, UP_* = 0.2, *RMSD*_R_ = 0.8, σ*_RMSD,R_* = 0.2, whereas in the one-up configuration, the overall means and standard deviations are *RMSD*_*L*_ = 0.7, σ_*RMSD, L*_ = 0.2, *RMSD*_*UP*_ = 1.3, σ*_RMSD, UP_* = 0.3, *RMSD*_R_ = 1.3, σ*_RMSD, R_* = 0.4. We do note the occurrence of a few higher deviation outliers in the all-down state, largely due to configurations of highly-flexible loops.

We also investigated spatial details of the fluctuations by presenting the RMSFs of each atom in the RBD. In the D-form (**Fig. S1 E**,**H**), the Up protomer demonstrates increased fluctuations in a large range of residues from 439 to 527, although the average magnitude of the fluctuations of VAL 483—the most highly-fluctuating residue—increase very little. Conversely, the fluctuations of L-down in the area decrease in magnitude across the board, while the fluctuations of R-down increase in the smaller area from about residues 478 to 487. Specifically, the average fluctuations of VAL 483 for the Up protomer go from 1.0 +/−0.1 to 1.2 +/−0.2; for the L-down go from 0.75 +/−0.05 to 0.54 +/−0.05; and for the R-down go from 0.62 +/−0.09 to 0.98 +/−0.06. It should be noted that these trends are largely preserved in the G-form **(Fig. S1B**,**D)**, although within error bars there is no increase in the R-down fluctuations. Specifically, the average fluctuations of VAL 483 for the Up protomer go from 1.0 +/−0.1 to 1.2 +/−0.2 (no change within error bars for the G-form); for the L-down go from 0.79 +/−0.05 to 0.42 +/−0.06; and for the R-down go from 0.7 +/−0.2 to 0.9 +/−0.1 (no change within error bars).

### Local effect of Glycosylation

Although the D614G substitution does not confer major global changes in exposure, small local changes near 614 occur due to the complex 2-antennae fucosylated glycan situated very close to the substitution site at residue 616. D614G leads to a reorientation of glycan 616, affecting the glycan coverage locally on the neighboring protomer FP. This local reorientation could possibly affect FP and even NTD targeting antibodies through extended glycan interactions (**Fig. S11**).

This glycan 616 can directly affect the shielding at the FP region. Glycans 282 and 603 from the same protomer are also located surrounding FP, contributing to the glycan coverage in this region. We observe that due to the increased flexibility around 614 resulting from the D to G change, the glycan at 616 can now sample a larger volume and interact with residues 46, 280 and 281 in the neighboring protomer in ∼24% of the ensemble (**Fig. S11C)**. Due to this reorientation of glycan 616 near the D614G, there is an overall increase in coverage for residues 826 to 855 in the all-down conformation (**Fig. S11C**).

## Supplementary Tables

**Table S1.** Average number of inter-protomer contacts between S1 and S2 regions.

| G-form all-down | G-form one-up | D-form all-down | D-form one-up |
| --- | --- | --- | --- |
| LU: $130 \pm 10$ | $120 \pm 10$ | $119 \pm 2$ | $143 \pm 7$ |
| UR: $130 \pm 10$ | $116 \pm 9$ | $123 \pm 9$ | $87 \pm 6$ |
| RL: $139 \pm 7$ | $127 \pm 10$ | $139 \pm 7$ | $127 \pm 10$ |

**Table S2.**
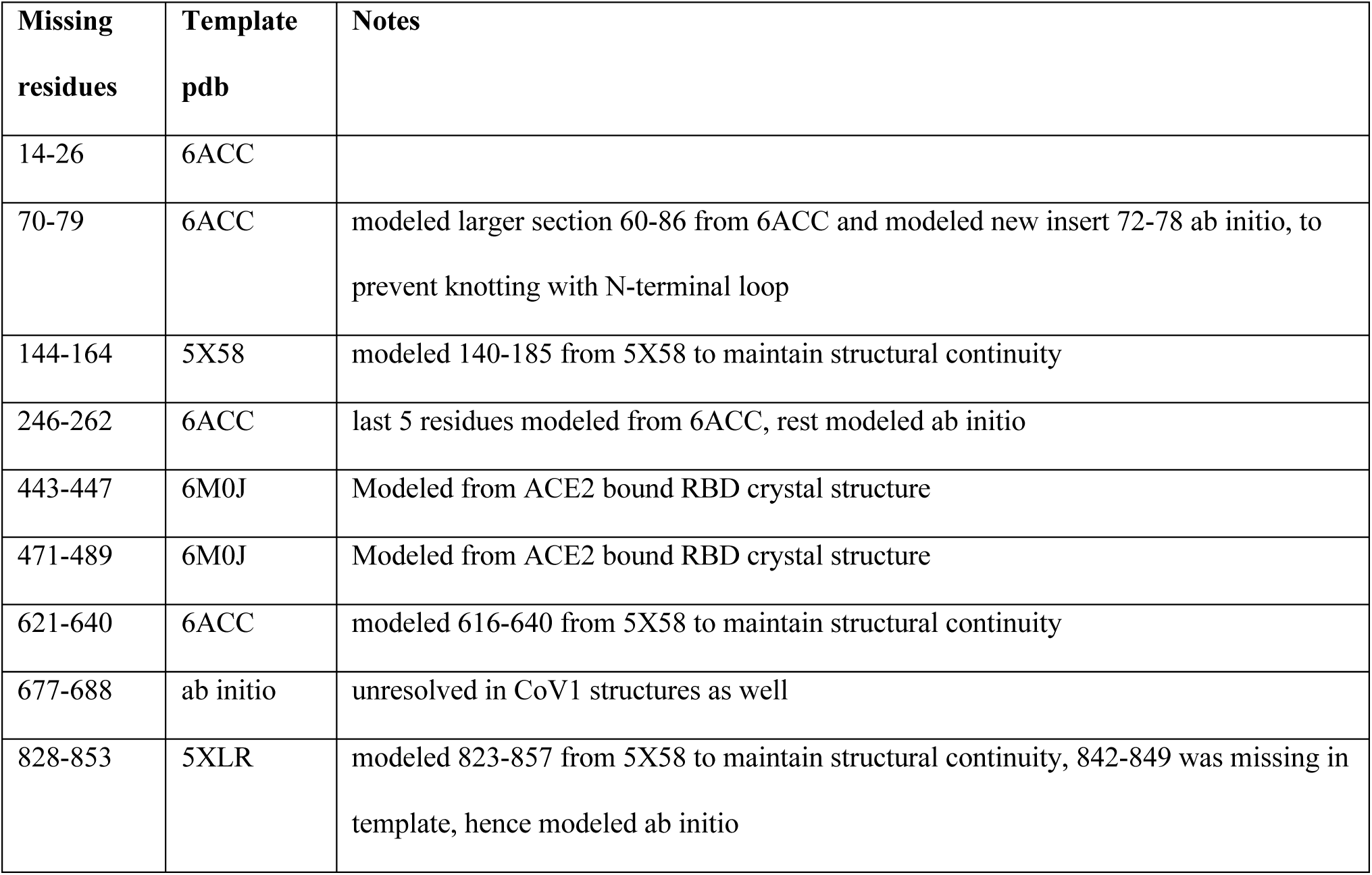
Template selection for SARS-COV2 Spike structural modeling

**Table S3.** SARS-CoV2 S glycoprotein N-glycan types. 19 of the 22 N-linked glycosylation sites are present in each of the three protomers of the model. Glycan type and constituent pyranose rings at each site correspond to that with highest relative population at each PNGS as observed in site-specific mass spectrometry results by Watanabe et al. (7)

| Spike ASN residue | Glycan Type | Constituent pyranose rings |
| --- | --- | --- |
| 17 | Fucosylated 3-antennae (FA3) | HexNAc(5)Hex(4)Fuc(1) |
| 61 | mannose-5 (M5) | HexNAc(2)Hex(5) |
| 74 | Fucosylated 3-antennae (FA3) | HexNAc(5)Hex(4)Fuc(1) |
| 122 | mannose-5 (M5) | HexNAc(2)Hex(5) |
| 149 | Fucosylated 2-antennae (FA2) | HexNAc(5)Hex(3)Fuc(1) |
| 165 | Fucosylated 3-antennae (FA3) | HexNAc(5)Hex(4)Fuc(1) |
| 234 | mannose-8 (M8) | HexNAc(2)Hex(8) |
| 282 | Fucosylated 3-antennae (FA3) | HexNAc(5)Hex(3)Fuc(1) |
| 331 | Fucosylated 2-antennae (FA2) | HexNAc(5)Hex(3)Fuc(1) |
| 343 | Fucosylated 2-antennae (FA2) | HexNAc(5)Hex(5)Fuc(1) |
| 603 | mannose-5 (M5) | HexNAc(2)Hex(5) |
| 616 | Fucosylated 2-antennae (FA2) | HexNAc(5)Hex(3)Fuc(1) |
| 657 | Fucosylated 2-antennae (FA2) | HexNAc(5)Hex(3)Fuc(1) |
| 709 | mannose-5 (M5) | HexNAc(2)Hex(5) |
| 717 | mannose-5 (M5) | HexNAc(2)Hex(5) |
| 801 | mannose-5 (M5) | HexNAc(2)Hex(5) |
| 1074 | mannose-5 (M5) | HexNAc(2)Hex(5) |
| 1098 | Hybrid | HexNAc(3)Hex(5) |
| 1134 | Fucosylated 2-antennae (FA2) | HexNAc(5)Hex(3)Fuc(1) |

## Supplementary Figures

**Fig. S1.**
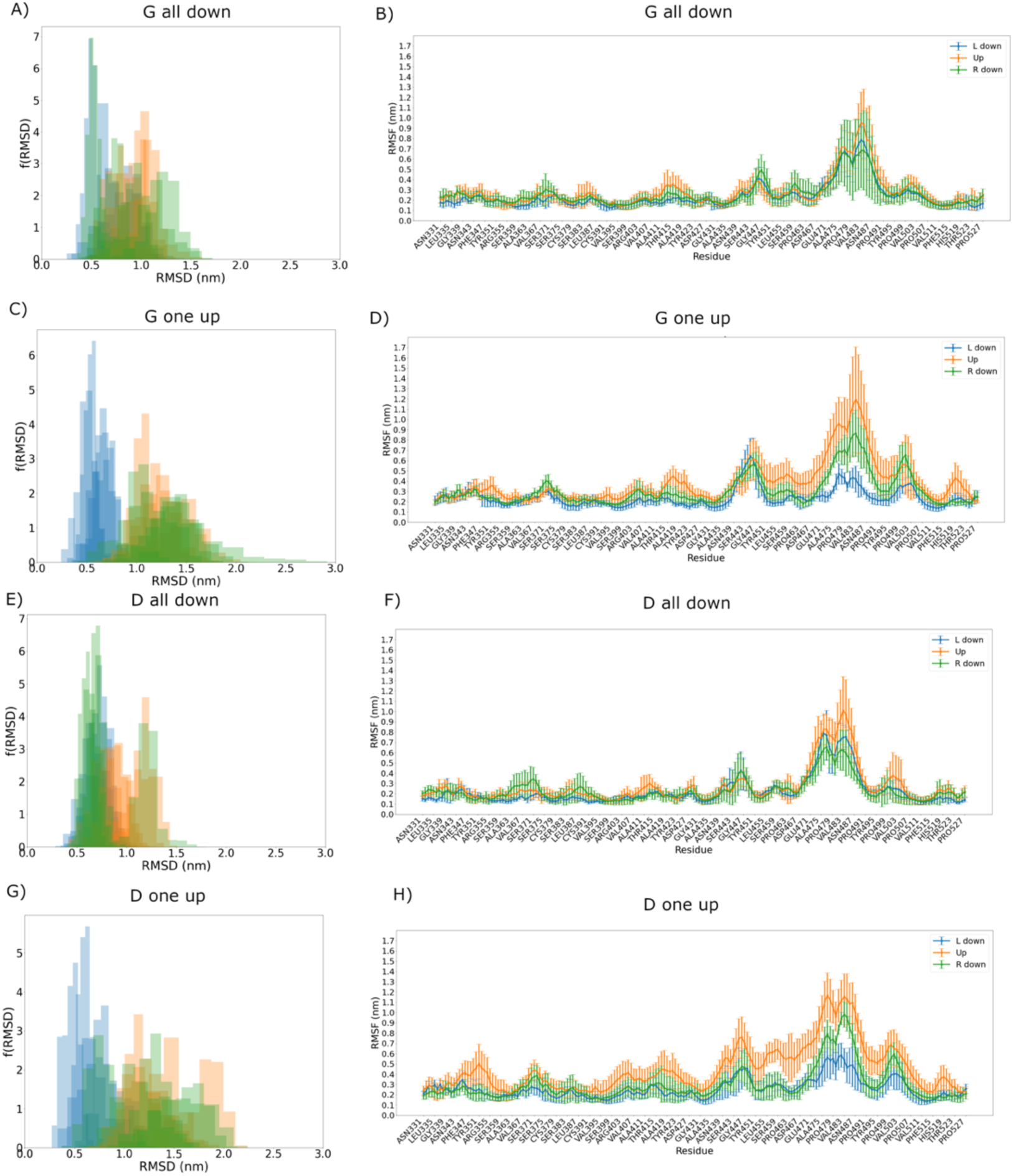
Root mean square deviations **(RMSDs)** and Root mean square fluctuations **(RMSFs) of the RBD.** **(A-D)** G-form, **(E-H)** D-form. **(A)/(C)** all-down configuration and **(B)/(D)** one-up configuration. **(A-B)** The distribution of RMSDs each replica is represented as a density histogram with twenty bins. Protomer L-down is blue; protomer Up is green; and protomer R-down is orange. **(C)-(D)** The fluctuation about its initial position of the Cα atom of each residue in the RBD. The thick lines represent the means; the error bars represent the standard deviations over five runs. **(E)/(G)** all-down configuration and **(F)/(H)** one-up configuration. **(E-F)** The distribution of RMSDs each replica is represented as a density histogram with twenty bins. Protomer L-down is blue; protomer Up is green; and protomer R-down is orange. **(G)-(H)** The fluctuation about its initial position of the C alpha atom of each residue in the RBD. The thick lines represent the means; the error bars represent the standard deviations over five runs. Figure rendered with the Matplotlib Python package version 3 (8).

**Fig. S2.**
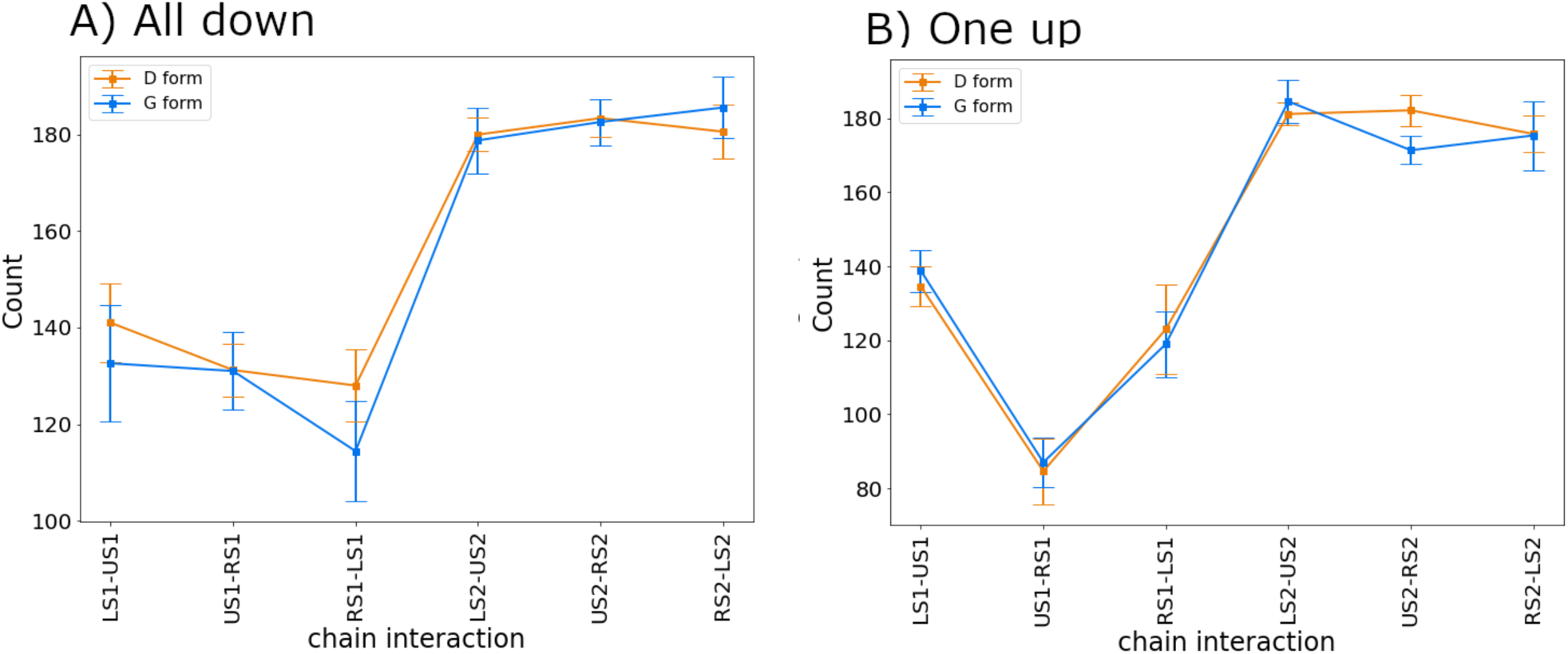
S1-S1 and S2-S2 contacts. Average total number of contacts at the S1-S1 and S2-S2 interfaces in **(A)** the all-down system and **(B)** the one-up system. For each set of simulations, error bars were calculated as standard error across five replicas. In the x-axis of all panels, ‘L’ denotes the L-down protomer, ‘U’ the Up protomer, and ‘R’ the R-down protomer. For instance, LS1-US1 represents the interactions between the S1 region of the L-down protomer and the S1 region of the Up protomer.

**Fig. S3.**
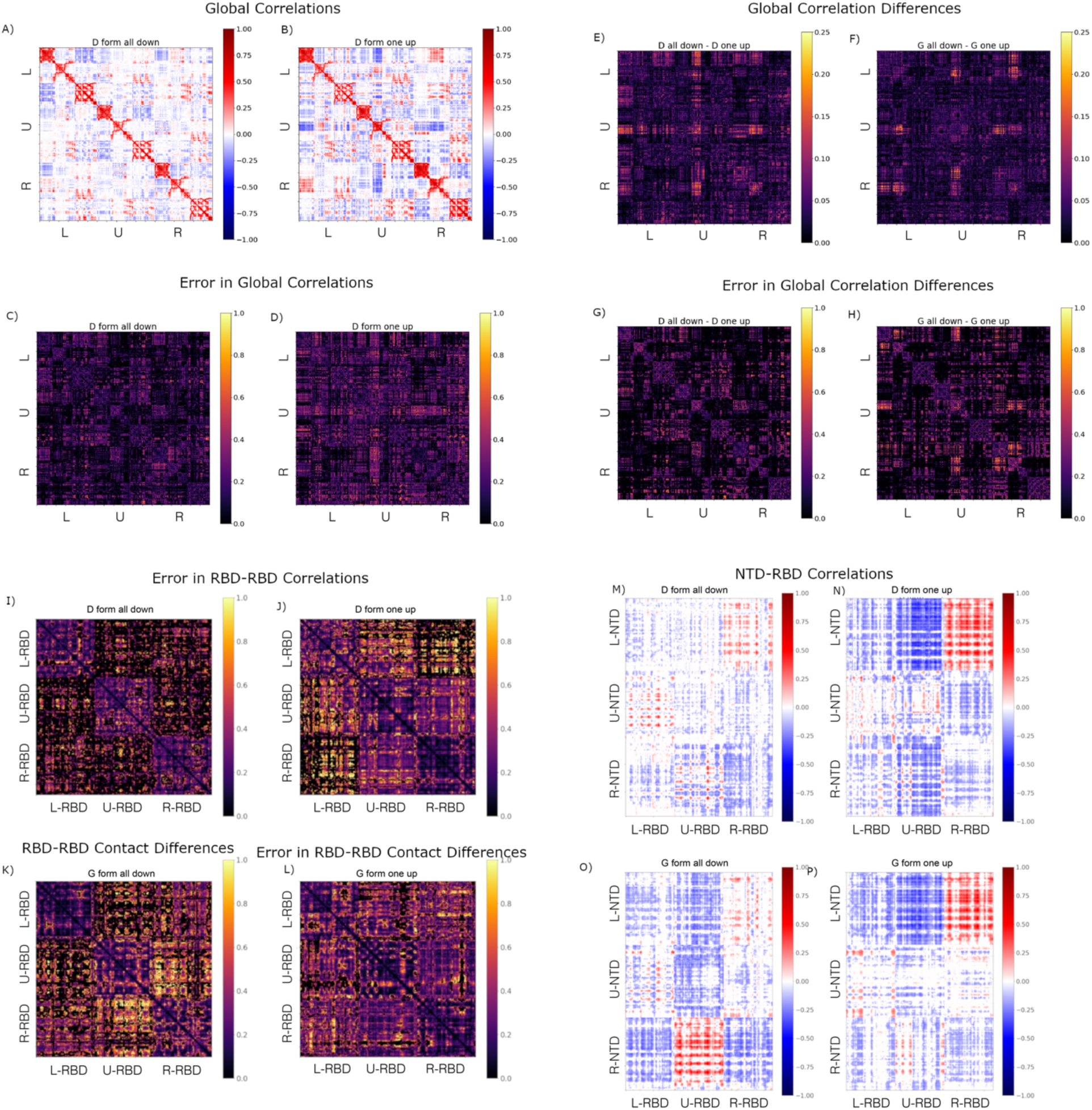
Global and local correlation matrices, differences, and fractional errors. **(A)-(B)** Full cross-correlation matrices of the D-form all-down **(A)** and D-form one-up **(B). (C)-(D)** Fractional errors in **(A)** and **(B)**, respectively, computed from the standard error across five replicas. **(E)-(F)** Differences between the matrices computed as 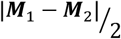, where ***M***_1_ is the first matrix mentioned in the title and ***M***_2_ is the second. **(H)-(G)** Fractional errors in **(E)** and **(F)**, respectively, computed from standard error propagation. **(I)-(L)** Fractional errors in RBD-RBD correlation matrix (**Fig. 3, S4**). **(M)-(P)** NTD-RBD correlation matrix. Red denotes positive and blue denotes negative cross-correlations in **A** and **M**. Higher the pixelation intensity, larger the magnitude of correlations. For the correlation difference and error plots, magnitudes increase from black (low) to yellow (high).

**Fig. S4.**
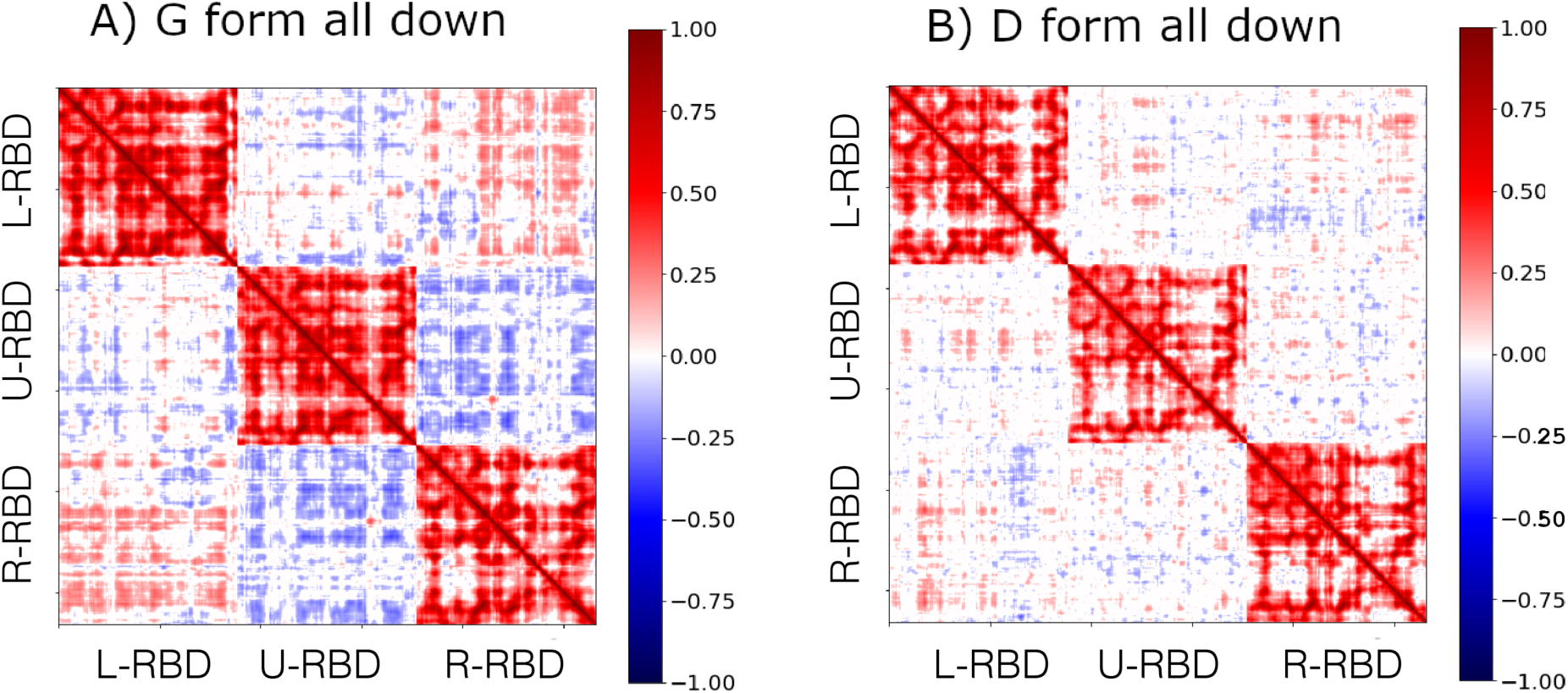
Residue-residue cross correlation matrices. **(A)** D-form all-down **(B)** G-form all-down. In the x and y axes, L-RBD, U-RBD, and R-RBD denote RBD regions of the L-down, Up, and R-down protomers, respectively. Red denotes positive and blue denotes negative cross-correlations. Higher the intensity of the pixels, stronger the magnitude of correlations. The diagonal blocks (top-left to bottom-right) denote high intra-domain correlation, and therefore have intense red pixelation. Inter-domain off-diagonal blocks have relatively stronger (though slightly asymmetric) correlations in G-form down (left) as compared to D-form down (right).

**Fig. S5.**
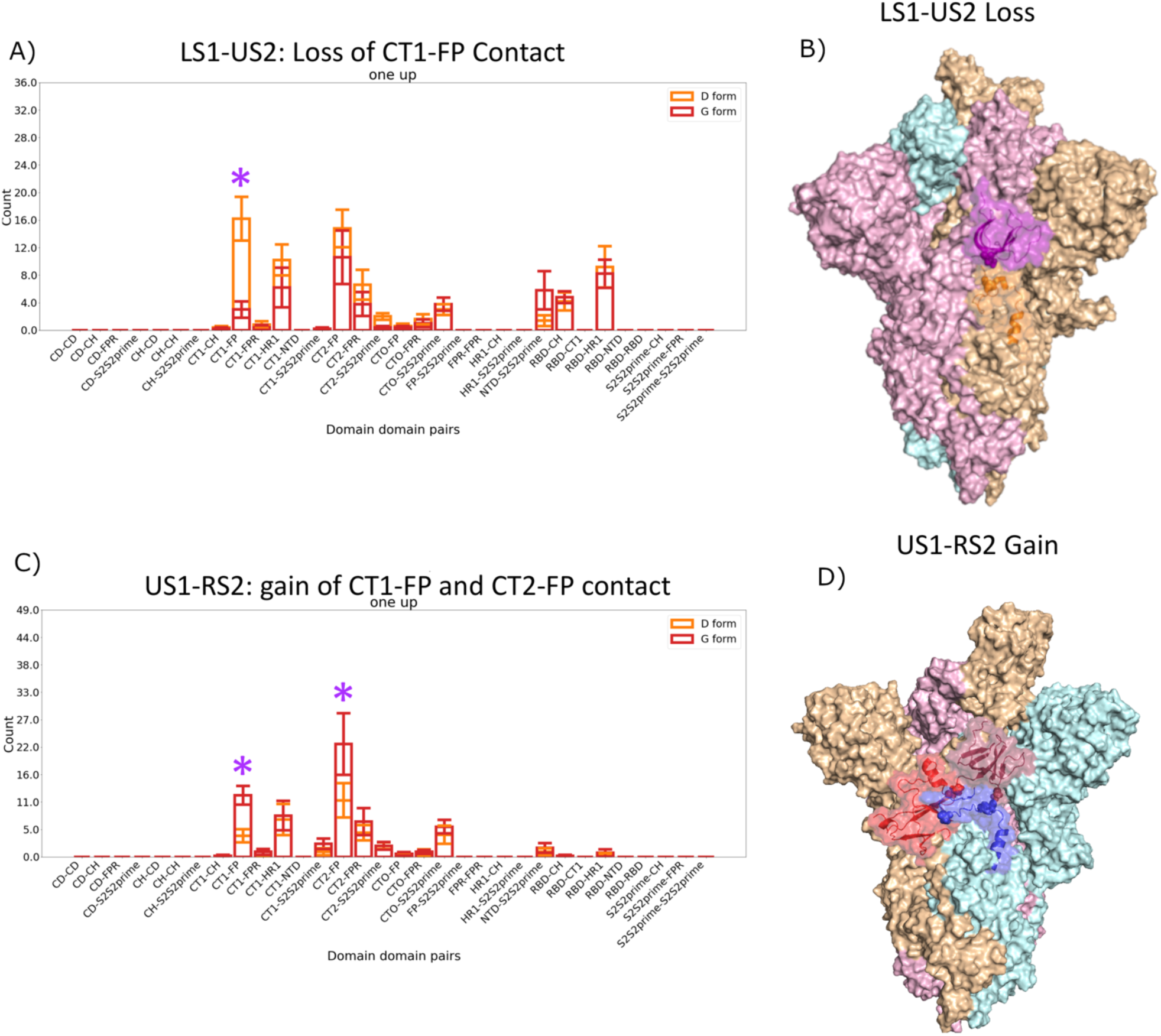
Domain-domain contact gain and loss for LS1-US2 and US1-RS2 interfaces. **(A)/(C)** Average number of contacts gained between forms by each system in terms of domain-domain interactions. Each bar represents the average number of contacts gained by the system in question between the two domains listed on the x axis, defined as the number of contacts in the D-form not in the G-form for D-form and the number of contacts in the G-form not in the D-form for the G-form. **(B)/(D)** Image of the two interfaces in question, with all three protomers shown as different colored surfaces, the domains involved in gain or loss shown as transparent surfaces with cartoon representations beneath, and the specific residues involved in the highest-frequency changing contact shown as spheres. Images rendered with PyMol (9).

**Fig. S6.**
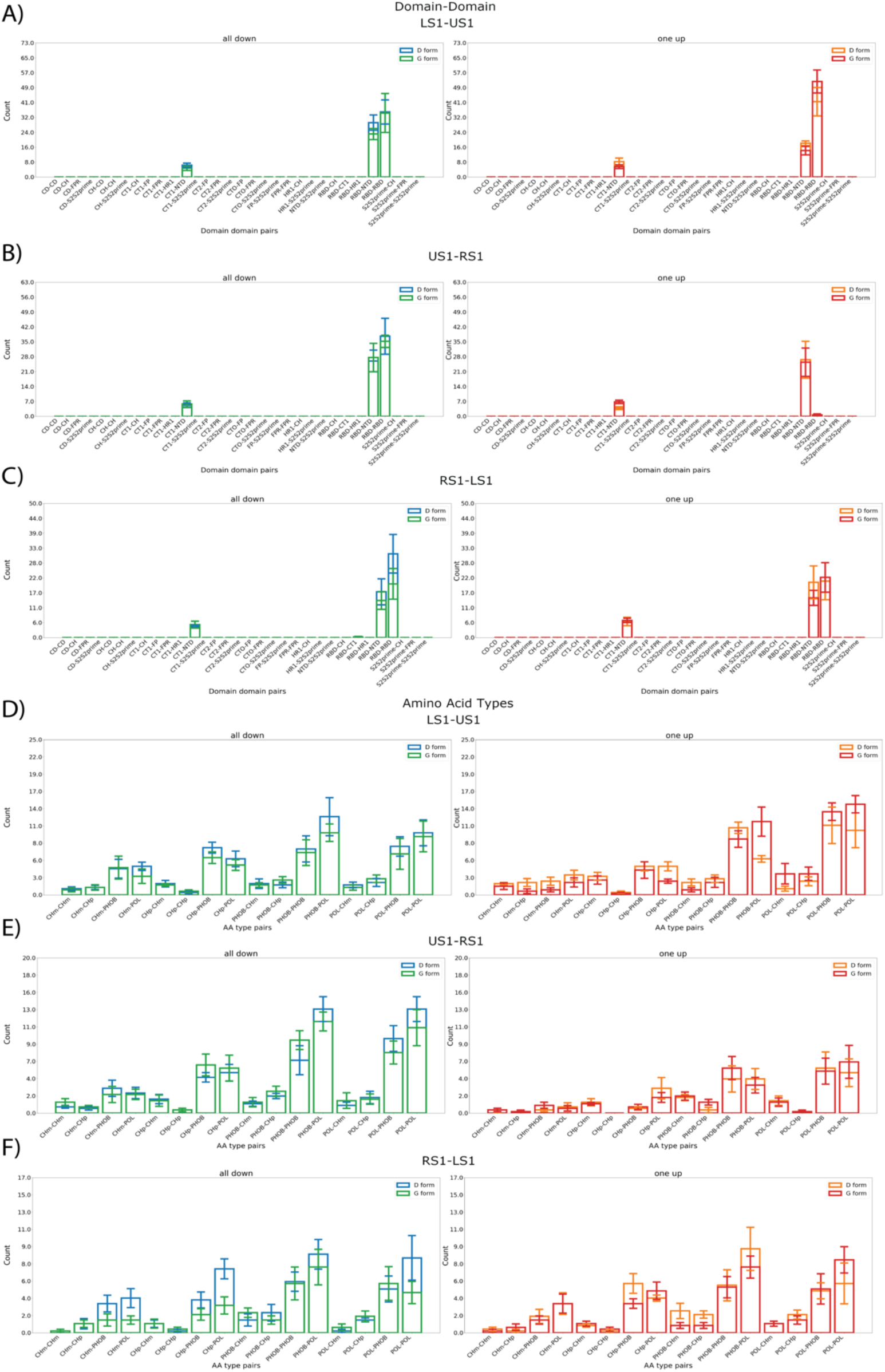
Details of changing inter-protomer contacts between S1 regions. **(A)-(C)** Number of contacts gained between forms by each system in terms of domain-domain interactions. Each bar represents the average number of contacts gained by the system in question between the two domains listed on the x axis, defined as the number of contacts in the D-form not in the G-form for D-form and the number of contacts in the G-form not in the D-form for the G-form. **(D)-(F)** Number of contacts gained between forms in terms of amino-acid type interactions. ‘Chm’ represents a negatively charged amino acid, ‘Chp’ represents a positively charged amino acid, ‘POL’ represents a polar amino acid and ‘PHOB’ represents a hydrophobic amino acid.

**Fig. S7.**
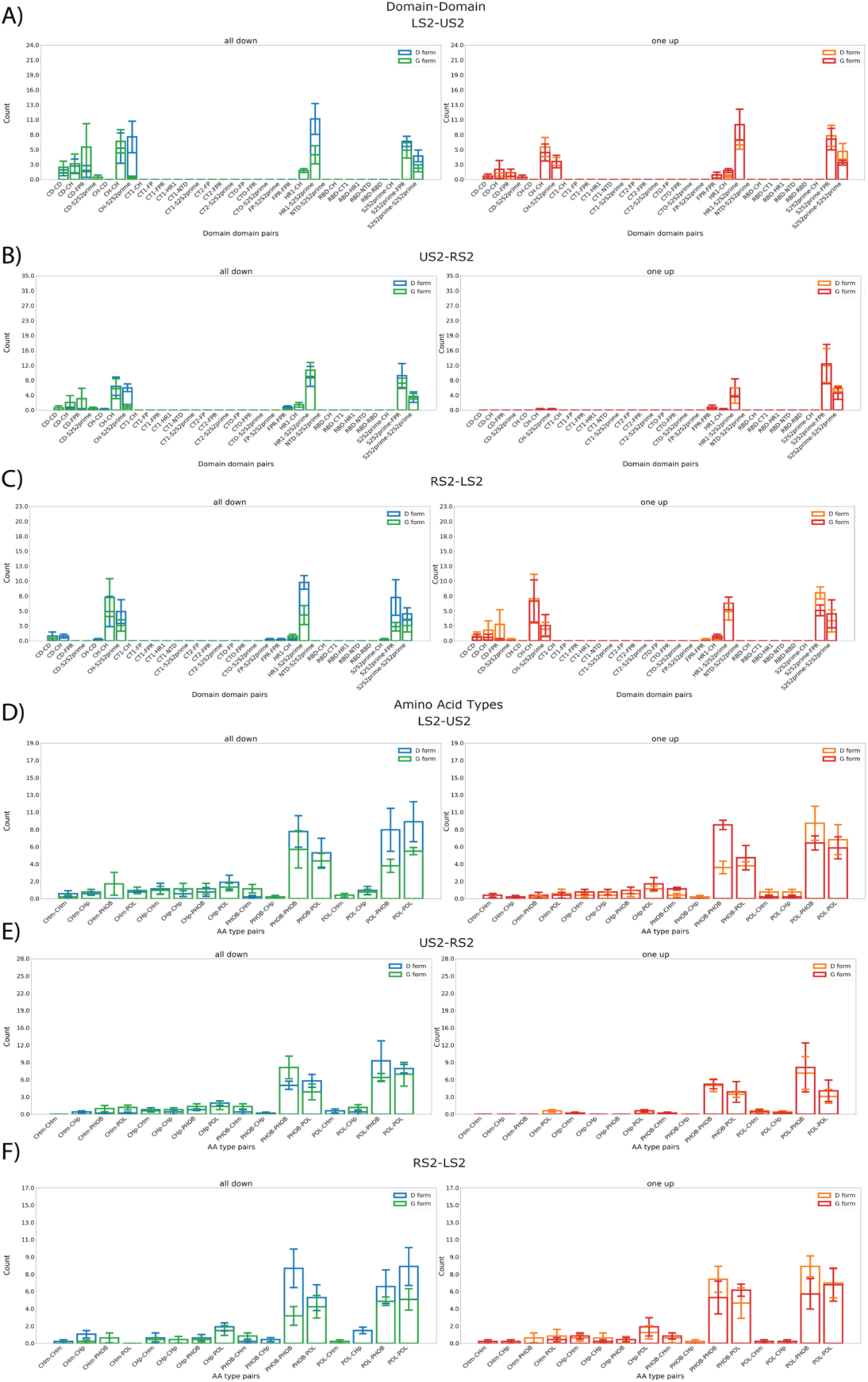
Details of changing inter-protomer contacts between S2 regions. **(A)-(C)** Number of contacts gained between forms by each system in terms of domain-domain interactions. Each bar represents the average number of contacts gained by the system in question between the two domains listed on the x axis, defined as the number of contacts in the D-form not in the G-form for D-form and the number of contacts in the G-form not in the D-form for the G-form. **(D)-(F)** Number of contacts gained between forms in terms of amino-acid type interactions. ‘Chm’ represents a negatively charged amino acid, ‘Chp’ represents a positively charged amino acid, ‘POL’ represents a polar amino acid and ‘PHOB’ represents a hydrophobic amino acid.

**Fig. S8.**
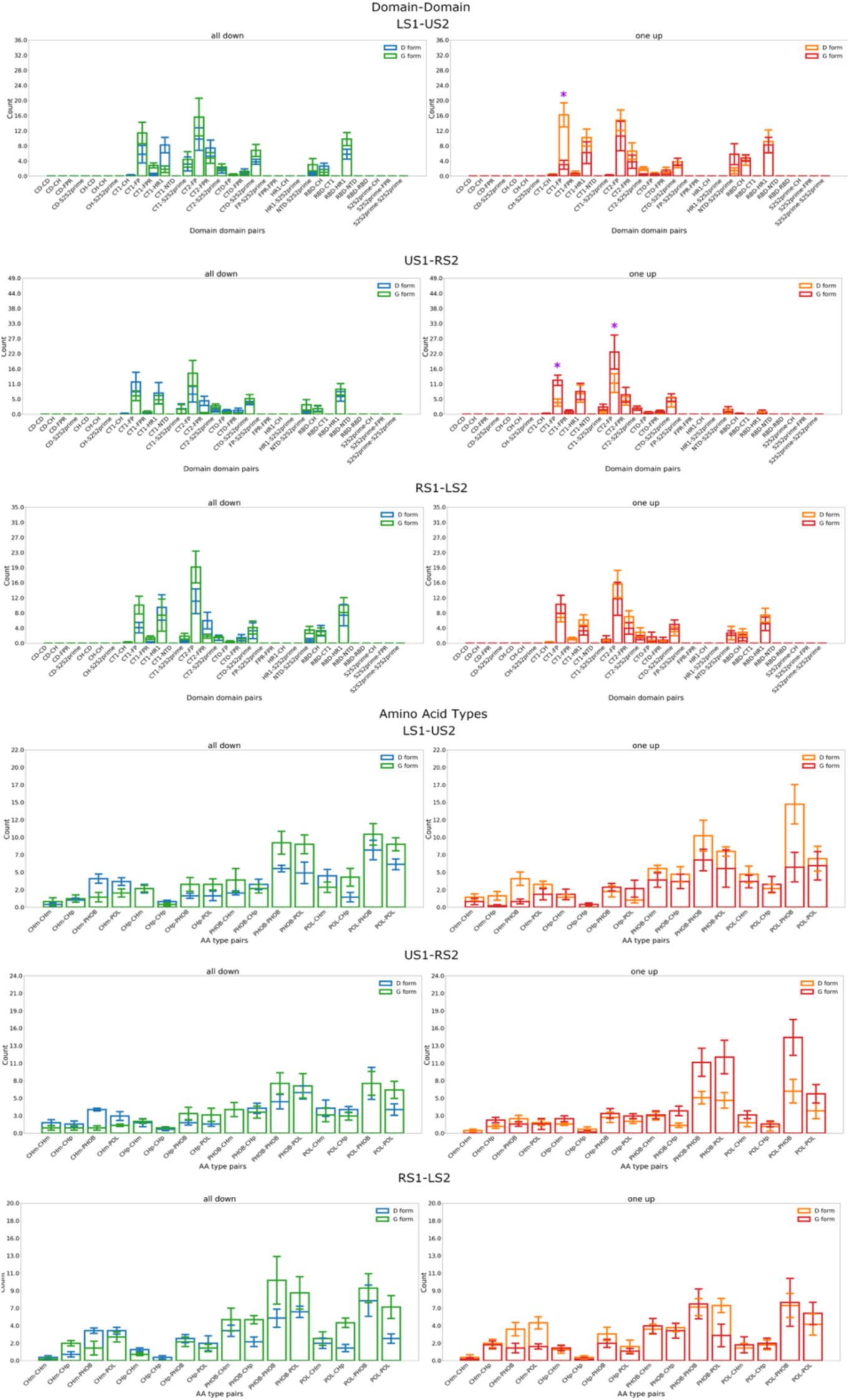
Details of changing inter-protomer contacts between S1 and S2 regions. **(A)-(C)** Number of contacts gained between forms by each system in terms of domain-domain interactions. Each bar represents the average number of contacts gained by the system in question between the two domains listed on the x axis, defined as the number of contacts in the D-form not in the G-form for D-form and the number of contacts in the G-form not in the D-form for the G-form. **(D)-(E)** Number of contacts gained between forms in terms of amino-acid type interactions. ‘Chm’ represents a negatively charged amino acid, ‘Chp’ represents a positively charged amino acid, ‘POL’ represents a polar amino acid and ‘PHOB’ represents a hydrophobic amino acid. Purple asterisks indicate domain-domain interactions of particular interest.

**Fig. S9.**
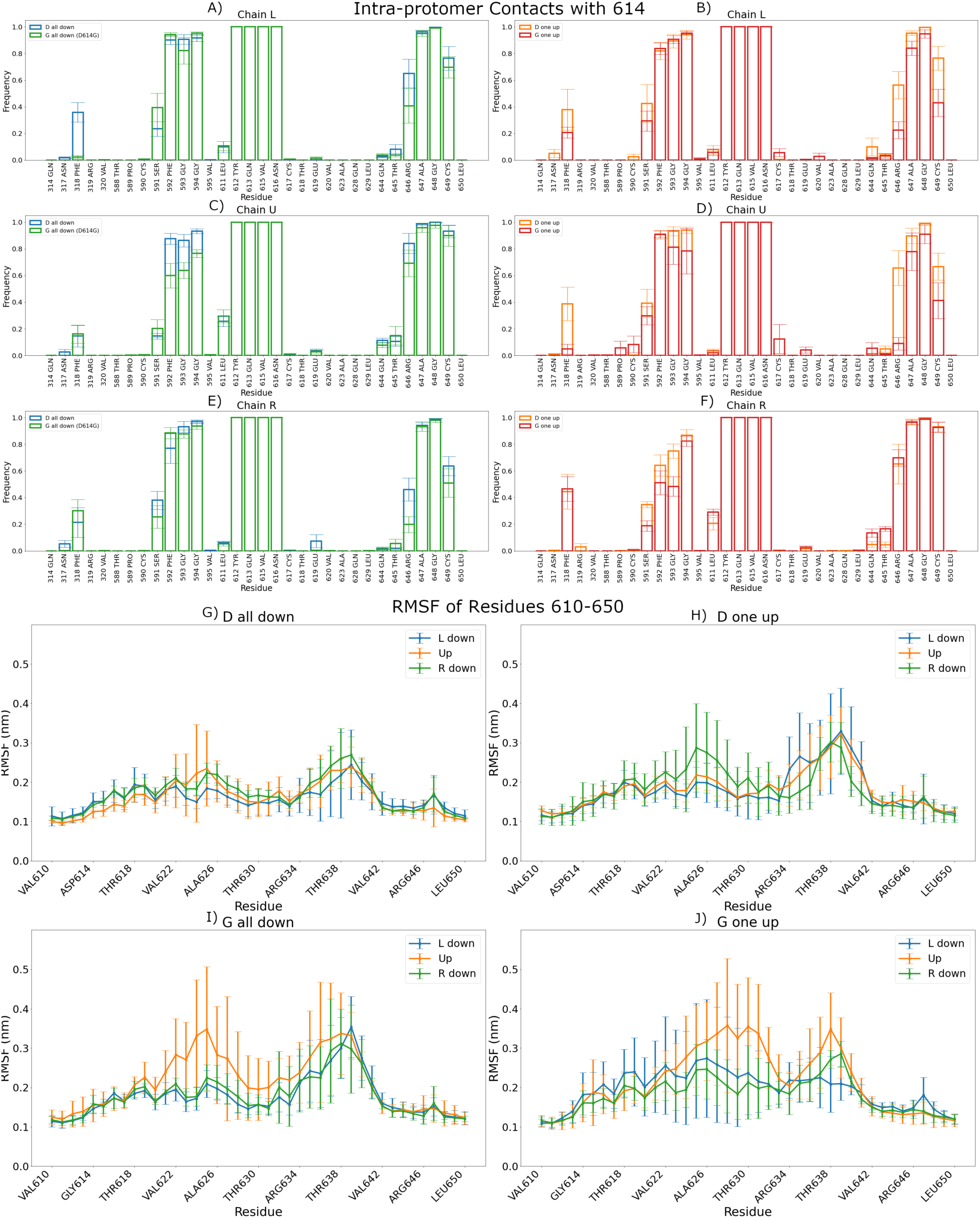
Increased flexibility with partial loss of intra-protomer hydrogen bond. **(A)-(F)** Intra-protomer contact frequencies with residue 614. **(G)-(J)** RMSFs of the region from residues 610-650 for D-form all-down **(G)**, D-form one-up **(H)**, G-form all-down **(I)**, and G-form one-up **(J)**. Error bars represent standard deviation over five replicas.

**Fig. S10.**
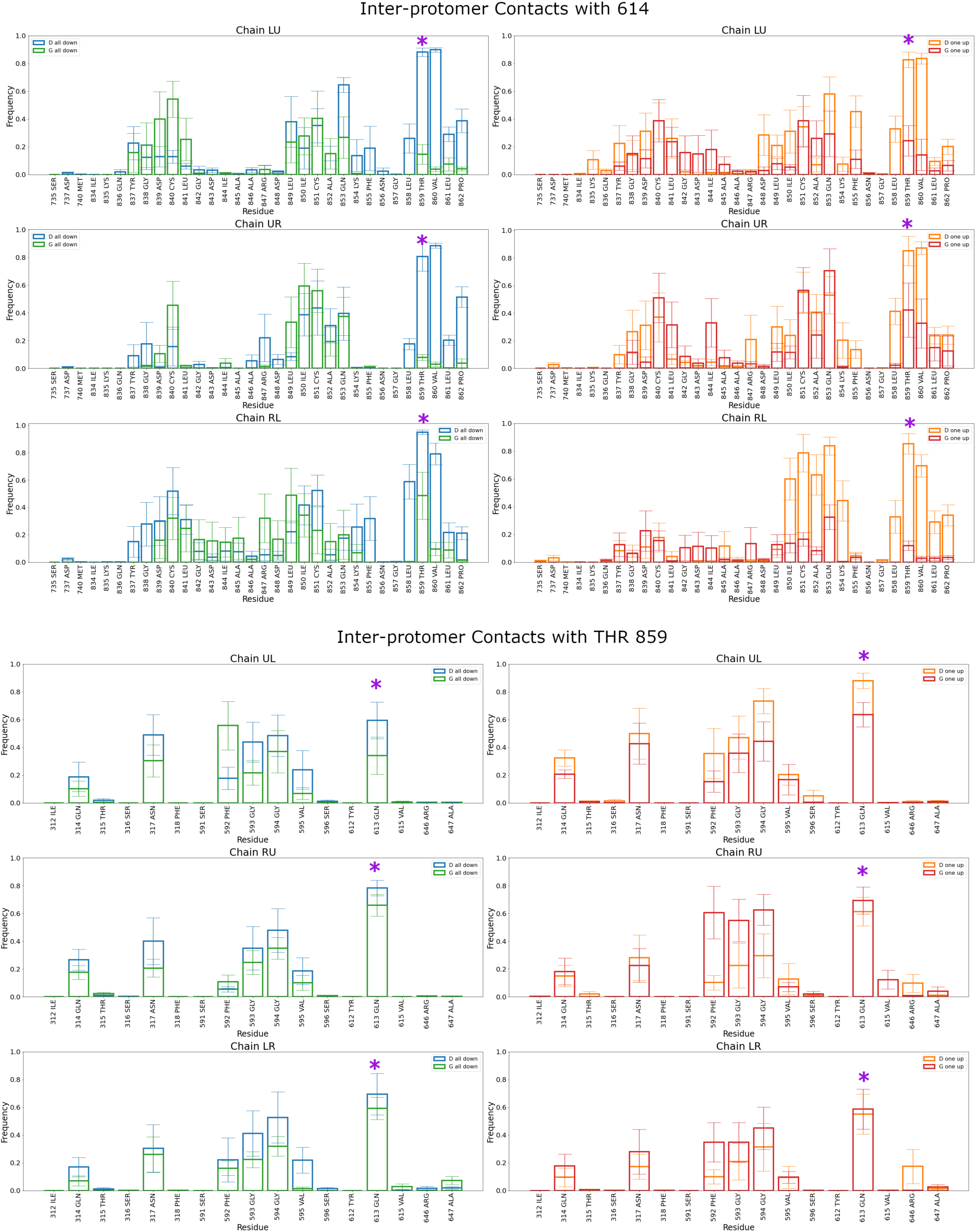
Inter-protomer contacts of interest. Inter-protomer contact frequency of **(Top)** residue 614 and **(Bottom)** residue THR 859. We show all contacts with non-zero contact frequency with the residues of interest between the different chains, except for THR 859 / residue 614 in **(Bottom)** since it is already shown in **(Top)**. Standard error computed across five different replicas. The purple asterisk on the top indicates the 614/859 contact. The purple asterisk on the bottom indicates the 613/859 contact.

**Fig. S11.**
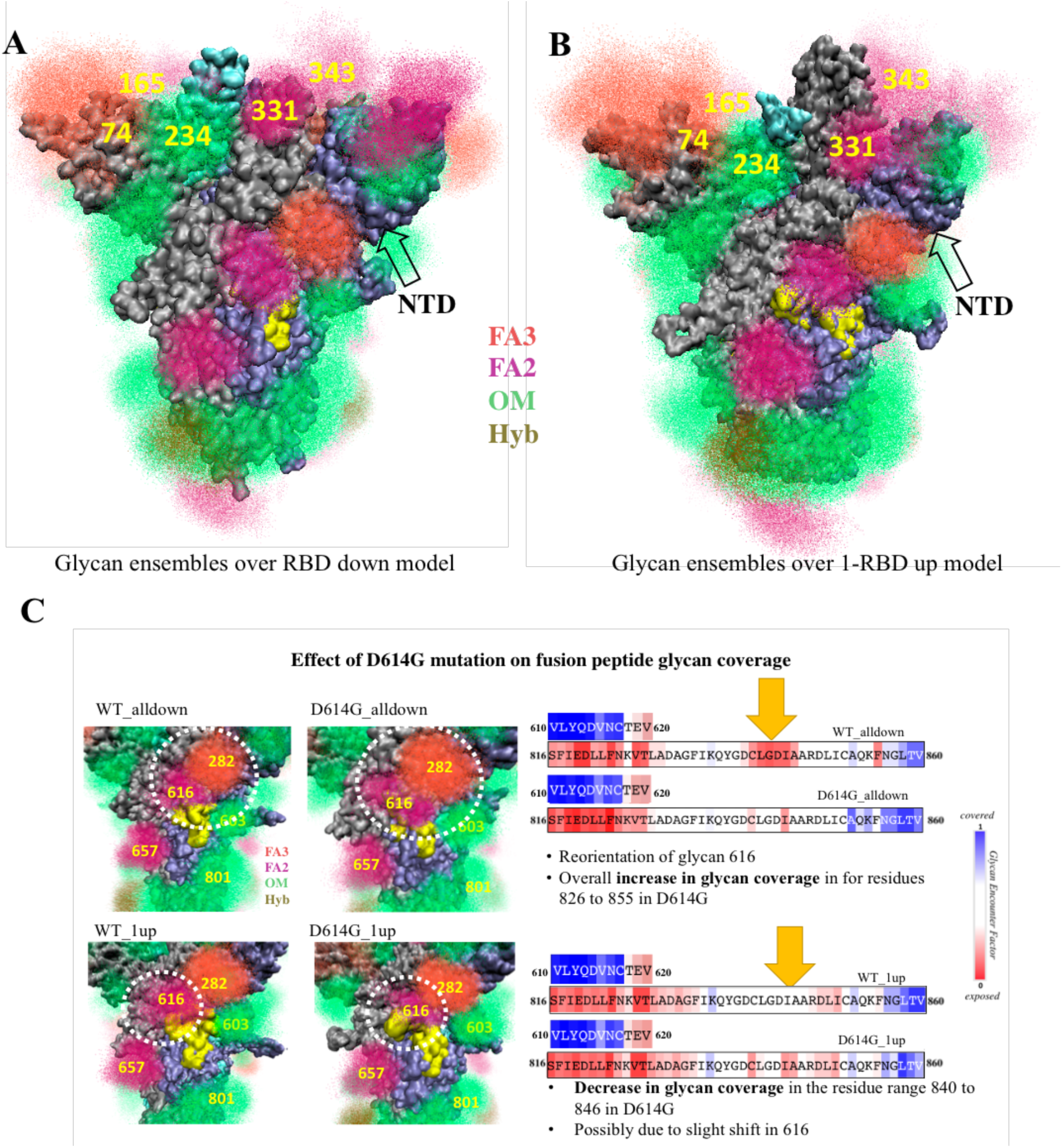
Potential immunological effects of SARS-CoV2 Spike protein glycosylation. (**A-B)**. Spike extracellular domain surface given in grey. The surface glycans are highly dynamic. Glycans overlaid from 250 different conformations are shown as points; this gives a “cloud” like rendition of the area most heavily occupied where the points are dense, and space that glycans sample less frequency. Fucosylated 2 and 3 antennae complex (FA2 and FA3) glycans, oligomannose (OM) glycans and hybrid (Hyb) glycans are colored according to key. (A) all-down and (B) one-up conformations shown. Glycans directly affecting RBD and NTD coverage change due to RBD opening are marked. Removal of glycan 165 has experimentally been shown to increase the population of protomers in the single-protomer up state. Removal of glycan 234 has experimentally been shown to decrease the population of protomers in the single-protomer up state (10). FP residue range 816 to 855 shown in yellow. (C) Effect of D614G substitution on FP coverage. N-glycans surrounding FP are numbered. Glycans at 282, 603 and 801 from the same protomer and glycan 616 from the neighboring protomer contribute to partial shielding of FP. On the left is the point cloud of glycans from 250 different conformations for the four different systems. White dashed circle shows the areas of change between D-form and G-form. Numbers label all glycans that touch the FP domain. On the right is the change in GEF superimposed on the relevant sequence.

**Fig. S12.**
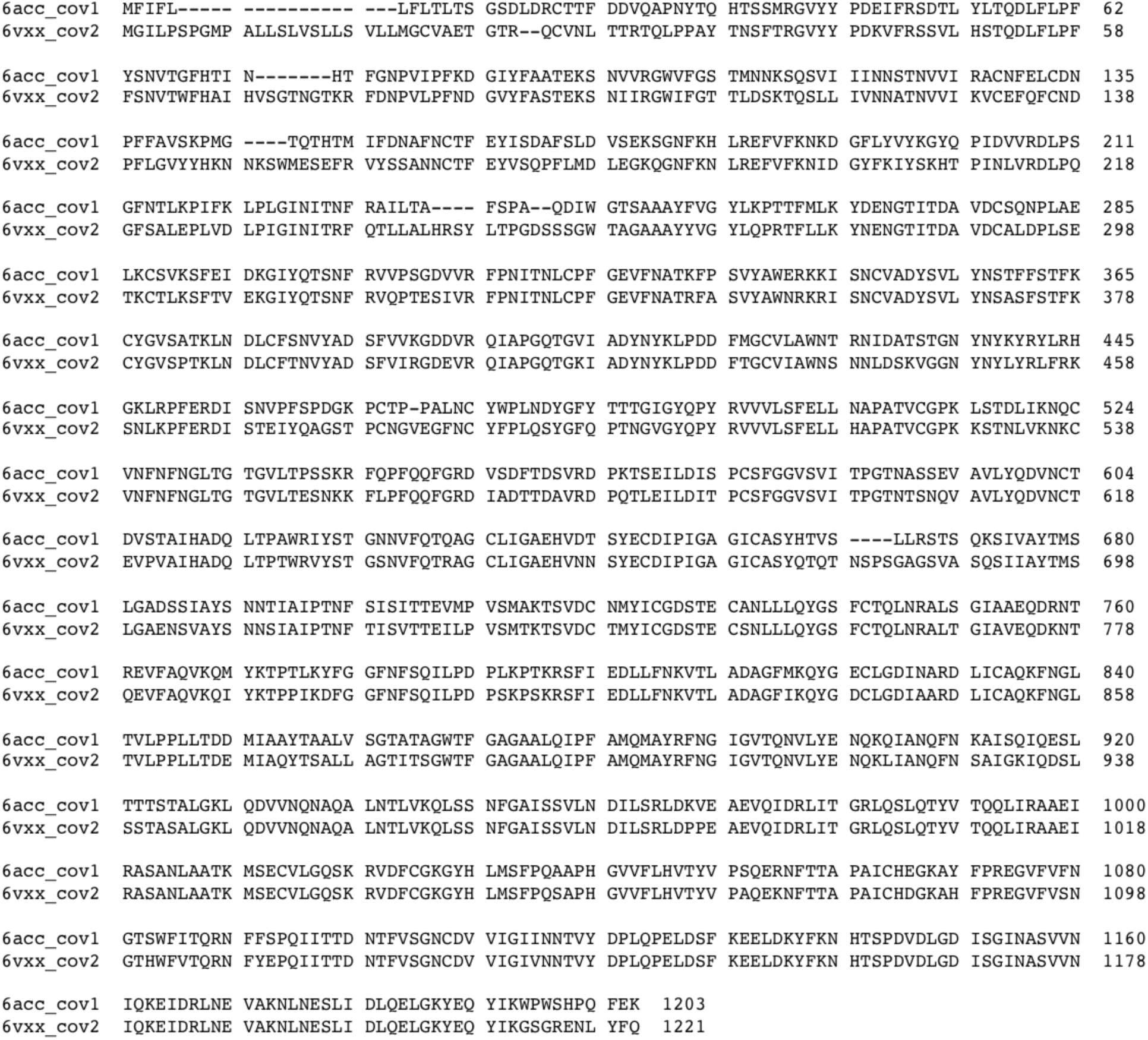
Structure based sequence alignment. Sequence alignment between SARS-CoV1 and SAR-CoV2 based on PDB structures 6ACC and 6VXX. Sequences given here are as found in the structures used.

## Ising Model of Covid Spike

Here we use a periodic three-spin Ising model to describe the occupation probability–and hence the population–of a particular thermodynamic macrostate through the Boltzmann distribution, which includes taking the exponential of the energy. It captures how a change in energies can lead to a large shift in the ratios of the respective populations.

Consider the one-dimensional periodic Ising model not in an external field, in which the Hamiltonian is defined as,

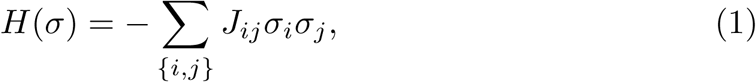

where the sum is across all nearest-neighbor pairs, including the pair {0, *N*} where *N* is the total number of spins.

We may think of the COVID Spike trimer in a highly simplified way as being a three-spin periodic Ising model, where each of the protomers is a “spin” that can take on an “up” state or a “down” state. Let us assume the following definitions for the interaction energies: *E*_*UU*_ is the magnitude of the interaction between a pair of protomers both in the up state, *E*_*DD*_ is the magnitude of the interaction between a pair of protomers both in the down state, *E*_*UD*_ is the magnitude of the interaction between protomers where the S1 region of the protomer in the up state interacts with the S2 region of the protomer in the down state, and *E*_*DU*_ is the magnitude of the interaction between protomers where the S2 region of the protomer in the up state interacts with the S1 region of the protomer in the down state. We do not assume these are equal since there is a known asymmetry in these interactions. Then the Hamiltonians of all microstates in the system are given as follows,

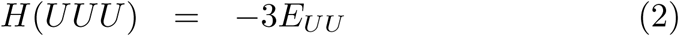

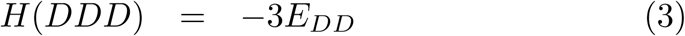

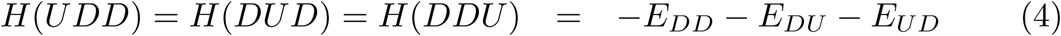

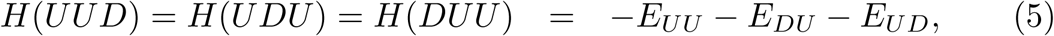

where we assume all interactions are overall favorable with negative interaction energies at body temperature, room temperature, etc.

The probability of being in a particular microstate is given by the Boltzmann distribution,

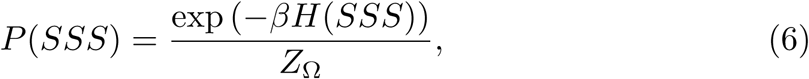

where *S* ∈{*U, D*}, *β* = (*k* _*B*_ *T*) ^−1^ is the Boltzmann factor, and *Z* _Ω_ is the normalizing constant or partition function.

Then let us consider the ratios of different macrostates to one another. We have four possible macrostates: all down (none up) (0 *U*), three (all) up (3 *U*), one up (1 *U*), and two up (2 *U*). Of these, all down and three up correspond to a single microstate, and one up and two up correspond to three microstates. Then the set of potentially interesting probability ratios is,

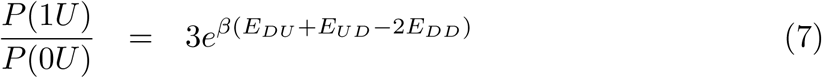

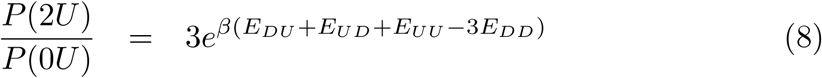

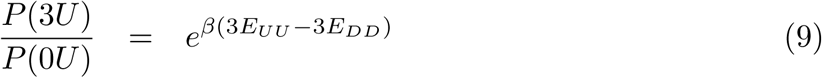

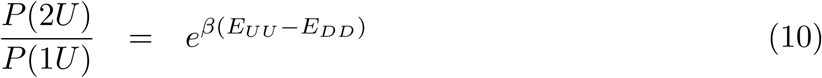

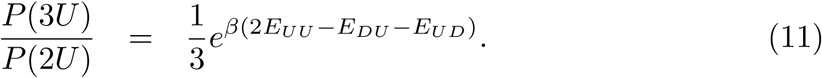

Assuming that most Cryo-EM and similar experiments have been performed at equilibrium, we note that there is a general lack of observation of the 2*U* and 3*U* states, which suggests that the difference between *E*_*UU*_ and *E*_*DD*_ is large with *E*_*UU*_ ≪ *E*_*DD*_, which would lead to a rapidly vanishing exponential in the comparison between 2*U* and 0*U*, 3*U* and 0*U*, and 2*U* and 1*U*, etc. Since most studies have seen an approximate 50/50 = 1 ratio of 1*U* to 0 *U* populations, this gives an order of magnitude estimate for the difference in *E*_*DD*_ and *E*_*DU*_ if we assume *E*_*DU*_ ≈ *E*_*UD*_, *T* = 310 K, and *k*_*b*_ = 1.38 × 10^−23^ m^2^kg s ^−2^K^−1^, of,

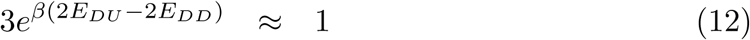

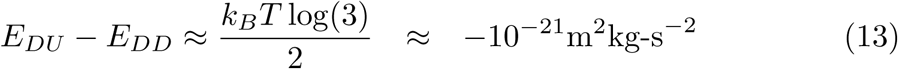

or approximately −0.006 eV, which is still a significant energy difference when considering the system as single particles in a highly idealized model.

If our results do point to an equalization of the energies between *E*_*DU*_ and *E*_*DD*_, then that would point to a shift in population from a 1:1 to a 3:1 ratio for 1*U* :0*U* as the argument of the exponential goes to zero and the exponential goes to one, meaning 75% 1*U* and only 25% 0 *U*, which could already be responsible for an increase in transmissibility.

It is unclear from our current results whether the mutation could also effect the *E*_*UU*_ interaction, but it is certainly not impossible. Since the Boltzmann distribution is an exponential, even relatively small changes in the argument can lead to large changes in the overall population of the states. A good avenue for future work would be to investigate the 2*U* and 3*U* states if possible.

One caveat to this interpretation is that it solely addresses thermodynamic contributions and does not consider the rates at which different states can transition from one to the other. If the rate of state transition is very low in comparison to the lifetime of the Spike between creation and ACE2 binding, then the thermodynamics are not the relevant quantity and what is more relevant is the initial configuration and–assuming that to be all down–the ease with which it can transition into the 1*U* conformation. This potential kinetic aspect of state transition would need to be addressed through different techniques, but it is also possible that the relaxation of the hydrogen bonding network induced by the mutant could have a significant impact on the kinetics as well as the thermodynamics.

